# Endogenous retroviruses are a source of enhancers with oncogenic potential in acute myeloid leukaemia

**DOI:** 10.1101/772954

**Authors:** Özgen Deniz, Mamataz Ahmed, Christopher D. Todd, Ana Rio-Machin, Mark A. Dawson, Miguel R. Branco

## Abstract

Acute myeloid leukemia (AML) is a highly aggressive hematopoietic malignancy, defined by a series of genetic and epigenetic alterations, which result in deregulation of transcriptional networks. One understudied but important source of transcriptional regulators are transposable elements (TEs), which are widespread throughout the human genome. Aberrant usage of these sequences could therefore contribute to oncogenic transcriptional circuits. However, the regulatory influence of TEs and their links to disease pathogenesis remain unexplored in AML. Using epigenomic data from AML primary samples and leukemia cell lines, we identified six endogenous retrovirus (ERV) families with AML-associated enhancer chromatin signatures that are enriched in binding of key regulators of hematopoiesis and AML pathogenesis. Using both CRISPR-mediated locus-specific genetic editing and simultaneous epigenetic silencing of multiple ERVs, we demonstrate that ERV deregulation directly alters the expression of adjacent genes in AML. Strikingly, deletion or epigenetic silencing of an ERV-derived enhancer suppressed cell growth by inducing apoptosis in leukemia cell lines. Our work reveals that ERVs are a previously unappreciated source of AML enhancers that have the potential to play key roles in leukemogenesis. We suggest that ERV activation provides an additional layer of gene regulation in AML that may be exploited by cancer cells to help drive tumour heterogeneity and evolution.

## Background

Acute myeloid leukaemia (AML) is characterized by clonal proliferation of immature myeloid cells. AML is highly heterogeneous at both the genetic and biological level, and individuals with AML accumulate a wide variety of genetic alterations that affect signalling pathways, transcription factors (TFs) and epigenetic modifiers^1^. In addition to genetic alterations, epigenetic processes have been shown to play key, and sometimes independent, dynamic roles in the molecular pathogenesis of AML^2,3^. For instance, altered chromatin landscapes, including DNA methylation^4^, histone modifications and chromatin accessibility^5,6^, are characteristics of AML subtypes. Genetic and epigenetic perturbations often target transcriptional regulatory networks, leading to dysregulation of transcriptional programs in AML and conferring a selective advantage^5,7^. During malignant transformation, leukaemia cells undergo continuous genetic and epigenetic diversification thereby increasing inter- and intra-patient tumour heterogeneity^3,8^, which directly reflects the complexity of leukemic transcriptional programs. One key component of transcriptional networks are transposable elements, which provide a rich source of tissue-specific *cis*-regulatory DNA sequences^9^. Despite extensive functional genomic analyses of AML, crucially the contribution of TEs to disease is currently unknown.

TEs have integrated into the human genome at different times throughout evolution and currently comprise around half of our genome. Based on their evolutionary origins, TEs vary with regards to their DNA structure. For instance, long terminal repeat (LTR) retrotransposons, which include endogenous retroviruses (ERVs), are composed of two LTRs that flank an internal retrovirus-derived coding region^10^. However, LTRs frequently recombine, leaving the majority of ERV elements as intact solitary LTRs, which contain functional *cis*-regulatory DNA sequences^11,12^. Therefore, ERVs are fixed in our genome but still maintain intrinsic regulatory capacity. Consistent with this, genome-wide assays have demonstrated that numerous LTR sequences carry hallmarks of active regulatory elements^13-20^. In a few instances, loss-of-function experiments have provided compelling evidence of LTR contribution to host gene regulation and cellular function in erythropoiesis^21^, innate immunity^18^, pregnancy^22^ and fertility^23^.

Various studies have documented widespread epigenetic and transcriptional deregulation of TEs in several cancer types, raising the possibility that TE-derived regulatory elements may be exploited to promote tumorigenesis^24,25^. Indeed, activation of LTR-based promoters initiates cancer-specific chimeric transcripts in Hodgkin lymphoma, melanoma and diffuse large B-cell lymphoma, amongst others^24,26,27^. However, studies to date have been centered on LTR promoter activity and their potential function as enhancers remains unexplored in human malignancies. Through the direct physical interactions with promoters, enhancers are especially important to regulate gene expression in a cell type-, temporal- and differentiation-stage-specific manner, all of which are essential for maintaining normal hematopoiesis. Indeed, dysregulation of specific enhancers, as well as global epigenetic disruption of the enhancer landscape have been shown to play critical roles in AML pathogenesis^28-30^. In this context, TEs are an ideal source of novel regulatory regions that could be co-opted in order to promote expression of genes essential for leukemic transformation and evolution in AML.

To explore the potential regulatory roles of TEs in AML, we utilized epigenomic and transcriptomic data from primary AML samples and leukemia cell lines. We have identified six ERV/LTR families with regulatory potential that harbor enhancer-specific epigenetic signatures and bind TFs that play key roles in haematopoiesis and in the pathogenesis of AML. Moreover, deletion of individual ERVs and epigenetic inactivation of an entire ERV family, demonstrate their direct roles in gene regulation. Strikingly, we found that either genetic or epigenetic perturbation of a single ERV-derived enhancer element leads to impaired cell growth by modulating expression of the *APOC1* gene, suggesting that the activation of this particular ERV has a driving role in leukemia cell phenotype.

## Results

### Identification of putative AML-specific regulatory TEs

To identify putative regulatory TEs, we generated DNase-seq data from three commonly used AML cell lines with different genetic and cytogenetic backgrounds: HL-60, MOLM-13 and OCI-AML3. Additionally, we analysed DNase-seq data from 32 AML samples generated by the Blueprint epigenome project^6^, and compared them with data from differentiated myeloid cells (macrophages, monocytes) from the same consortium (Figure 1A). We overlapped DNase hypersensitive sites (DHSs) with the complete Repeatmasker annotation and compared the DHS frequency at each repeat family with random controls (Supplementary Table 1). We identified twelve repeat families that were enriched for DHS-associated copies in at least one of the AML cell lines and in 10% or more of the AML samples (Figure 1B). Five of these repeat families (three of which are not TEs) were highly enriched across all samples, including macrophages and monocytes, as well as mobilized CD34+ cells (data from the Roadmap epigenomics project), suggesting little cell specificity. The remaining seven families displayed more variability between AML samples and, notably, tended to display little to no enrichment in differentiated myeloid cells (Figure 1B). Nearly all families were also DHS-enriched in CD34+ cells, suggesting an association with a stem cell state, which may be exploited by cancer cells to promote cell proliferation and survival. In contrast, the DHS enrichment of LTR2B elements appeared to be AML-specific and therefore associated only with the disease state. Analysis of an independent dataset of 32 AML samples from the Bonifer lab^5^ confirmed the DHS enrichment at all of the above families, and identified additional weaker associations, including with several *Alu* subfamilies (Supplementary Figure 1A). For stringency, we focused on families that were DHS-enriched in both datasets, all of which are LTRs from ERVs: LTR2B, LTR2C, LTR5B, LTR5_Hs, LTR12C, LTR13A. We excluded the internal portion of HERVK (HERVK-int) because its enrichment was largely due to its LTRs (LTR5B, LTR5_Hs; Supplementary Figure 1B). We will collectively refer to the six selected ERV families as ‘AML DHS-associated repeats’ (A-DARs). The oldest A-DARs (LTR5B, LTR13A) date back to the common ancestor between hominoids and old world monkeys, whereas the youngest (LTR5_Hs) are human-specific^31^.

**Figure 1.**
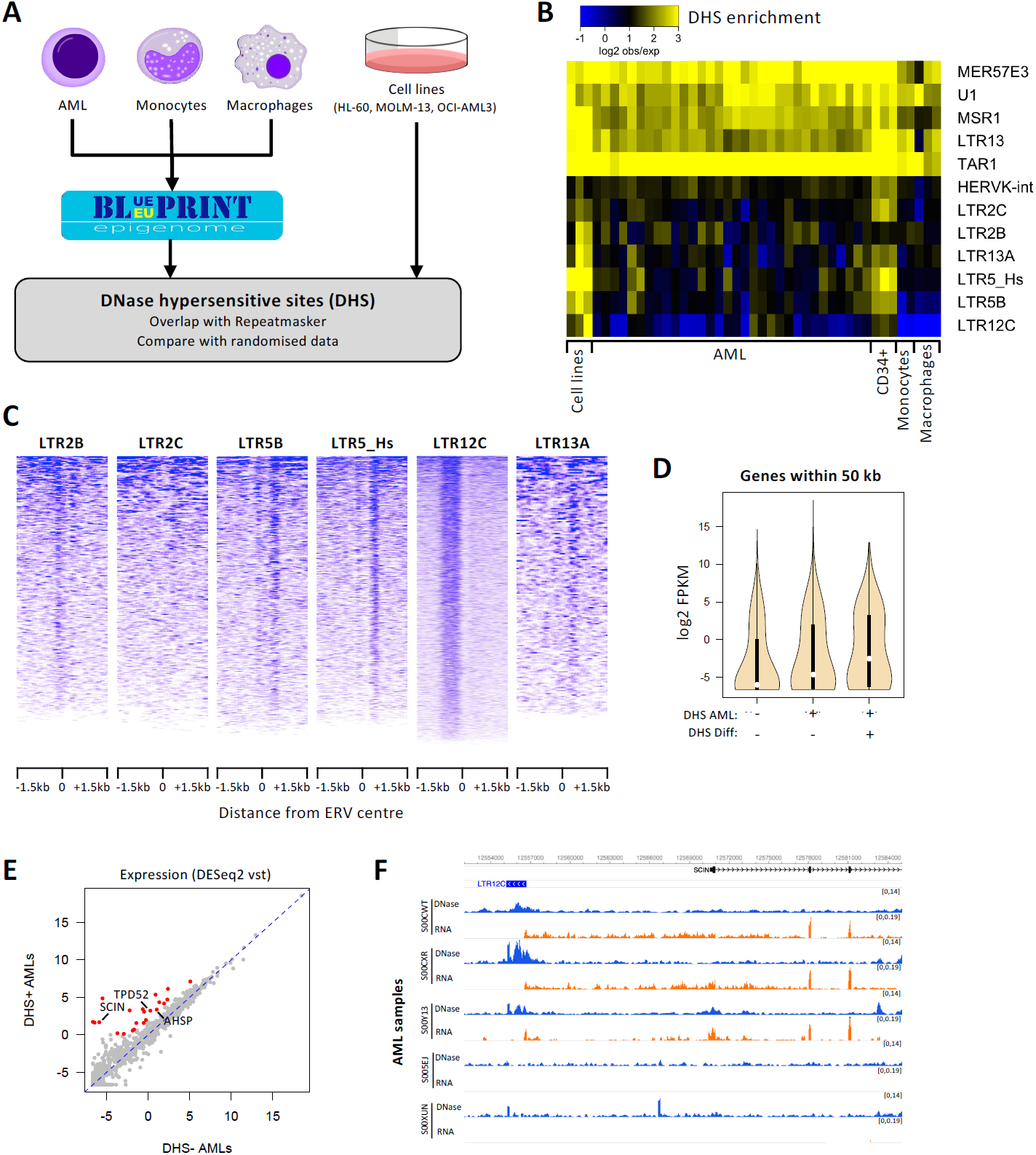
ERVs with regulatory potential are activated in AML and associated with gene expression differences (hematopoietic cells credit: A. Rad and M. Häggström; CC-BY-SA-3.0 licence). **A.** Schematic of the strategy to detect repeat families associated with open chromatin in AML. **B.** Heatmap of the observed/expected enrichment for DHSs in selected repeat families. Cell lines are presented in the following order: HL-60, MOLM-13, OCI-AML3. **C.** DNase-seq profile across all elements of each AML DHS-associated repeat (A-DAR) families in OCI-AML3. **D.** Gene expression average across all Blueprint AML samples for genes within 50 kb of A-DARs with or without a DHS in AML and/or in differentiated cells. **E.** For each gene lying near an A-DAR element, we compared its expression in AML samples where the respective ERV has a DHS, versus AML samples where the DHS is absent. Expression values were normalized using the variance stabilizing transformation (vst; log2 scale) in DESeq2. Highlighted are genes with >4-fold difference and vst>0. **F.** Example of a gene (*SCIN*) that displays a strict correlation between its expression (orange) and the presence of a DHS peak (blue) at a nearby LTR12C element in different AML samples.

The DNase-seq profiles across each ERV displayed a consistent pattern for elements of the same family in AML cell lines (less evident for LTR2C), suggestive of TF binding events within these ERVs (Figure 1C displays OCI-AML3 profiles). This pattern was also notable in primary AML cells, albeit variable between samples (Supplementary Figure 1C), reflecting the heterogeneity of this disease. Out of a total of 4,811 A-DAR elements, 80-661 (median 263) overlapped a DHS in AML samples from the Blueprint dataset and 223-1,349 (median 508) in the Assi et al dataset. As heterogeneity in AML is partly driven by genetics, we hypothesized that variation in DHS frequency at A-DARs could reflect distinct mutational profiles. To test this, we measured inter-sample correlations in the DHS patterns of A-DARs, which revealed distinct clusters associated to the mutational profile in AML patient samples (Supplementary Figure 2A). Although there was no strict association with particular AML subtypes, we found that samples with *NPM1* mutations were better inter-correlated than those without (Supplementary Figure 2B). The same was true for samples with *FLT3*-ITD and *DNMT3A* mutations, which frequently co-occur with *NPM1* mutations, as well as those with *CEBPA* mutations (Supplementary Figure 2B). Specific mutations may therefore contribute to ERV activation in AML, although other characteristics of the malignancy are also likely to affect them.

**Figure 2.**
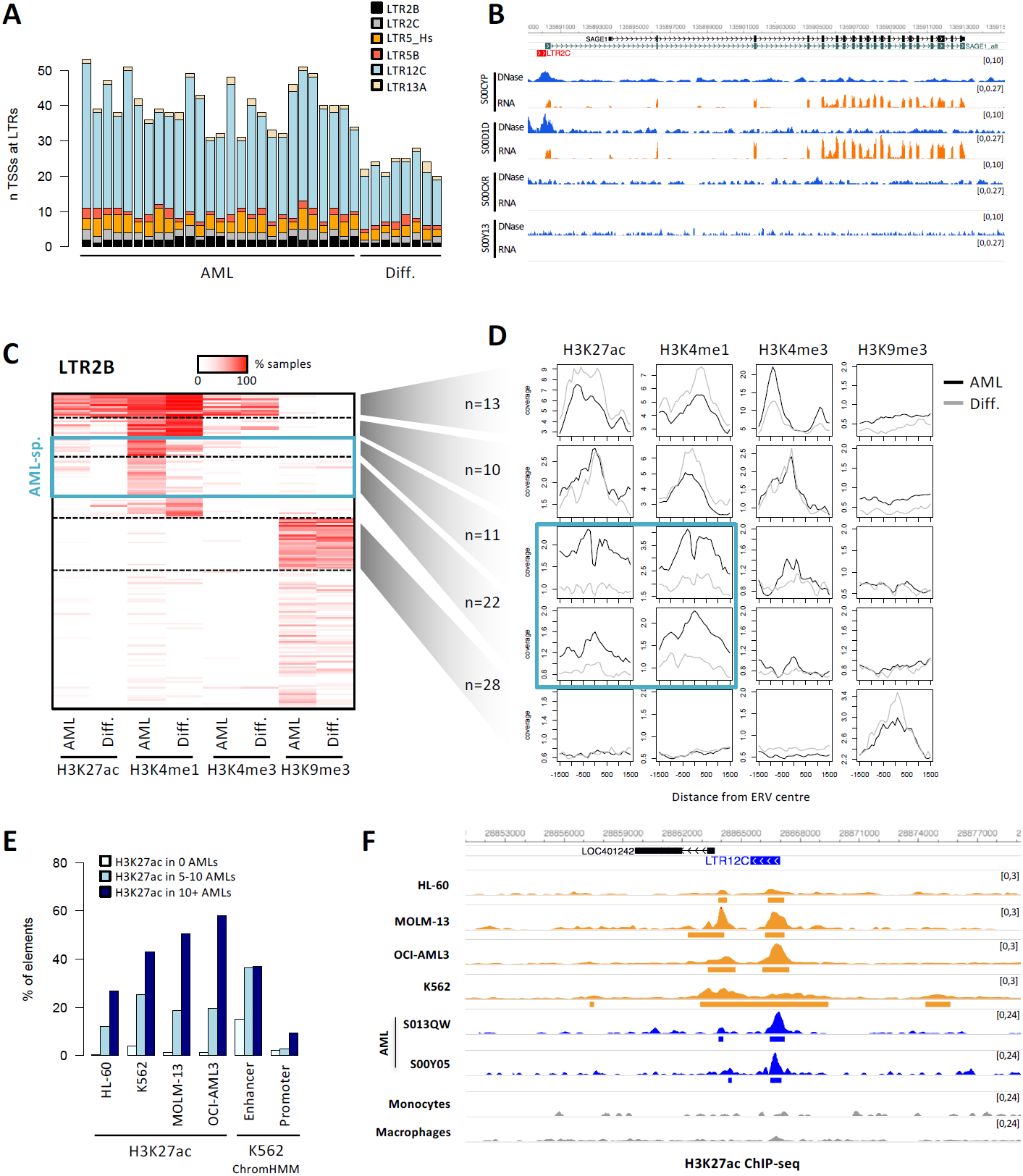
A-DARs bear signatures of enhancer elements. **A.** Number of transcriptional start sites of spliced transcripts that overlap with A-DAR elements in AML or differentiated myeloid cells. **B.** Example of a LTR12C element that generates an alternative promoter that drives the expression of *SAGE1* in AML samples where this element is active. **C.** Heatmap of overlap between LTR2B elements and histone modification peaks. Colour intensity represents the percentage of AML or differentiated cell samples where overlap is observed. Dashed lines segregate clusters identified by k-means clustering. **D.** Average ChIP-seq profiles for LTR2B elements within specific clusters define in C. Blue boxes highlight two clusters where H3K4me1 and H3K27ac levels are higher in AML compared with differentiated cells. **E.** Percentage of A-DAR elements that overlap H3K27ac peaks in different cell lines, or that are classified as enhancer or promoters in ChromHMM data from K562 cells. A-DAR elements were subdivided according to the number of AML samples displaying overlap with H3K27ac. **F.** Example of a LTR13A element where three cell lines reproduce the AML-specific H3K27ac marking observed in AML samples. Peaks called by MACS2 are depicted underneath each track.

### A-DAR chromatin status correlates with nearby gene expression

To test whether A-DARs were associated with gene activation, we analysed matching DNase-seq and RNA-seq data from the Blueprint consortium (n samples: 27 AML, 6 macrophage, 8 monocyte). ERVs can affect the expression of proximal genes, but also act at a distance via long-range interactions in 3D space^19,32^. However, long-range interactions display substantial cell specificity, namely within the hematopoietic system^33^. Given the heterogeneity between AML samples and the lack of matching Hi-C data, we stringently focused our analysis on genes within 50 kb of an ERV from the selected families. Genes close to A-DAR elements with DHS in two or more AML samples displayed higher expression levels than those close to A-DAR elements without DHS (Figure 1D). This was more pronounced for ERVs with DHS also present in differentiated cells. Even though such bulk correlations are only suggestive of a regulatory role of ERVs, we found individual elements with strong supporting evidence for their regulatory activity, as the expression levels of their adjacent genes were >4-fold higher in AML samples with DHS at a given ERV, versus those without (Figure 1E; see also Supplementary Table 6). This included a strict correlation between chromatin accessibility at a LTR12C element and the expression of the *SCIN* gene (Figure 1F). Notably, low *SCIN* expression is associated with an adverse AML prognosis^34^. Two other genes of interest for which expression also correlates with a DHS at nearby ERVs are *TPD52* and *AHSP*, whose overexpression in AML is predictive of poor and favorable outcomes, respectively^35,36^. These data suggest that at least some A-DAR elements gain gene regulatory activity in AML, which correlates with disease outcomes.

### A-DARs bear the chromatin signatures of enhancer elements

DNase hypersensitivity is associated with both active gene promoters and distal enhancers. LTR12C elements, for example, were previously shown to frequently act as alternative gene promoters in different cell types, including hepatocellular carcinoma^37^ and cell lines treated with DNMT and HDAC inhibitors^38^. In contrast, LTR5_Hs (HERV-K) elements appear to mainly act as distal enhancer elements in embryonic carcinoma cells and embryonic stem cells^19,20^. We therefore aimed to establish whether A-DARs could act as promoters and/or enhancers in AML.

To test for gene promoter activity, we performed *de novo* transcriptome assembly in AML samples and differentiated myeloid cells, and calculated the number of spliced transcripts for which the transcriptional start site (TSS) overlapped an A-DAR element. AML samples displayed 31-53 such transcripts, whereas differentiated cells had 20-28, most of which emanated from LTR12C elements (Figure 2A).

We identified 82 spliced transcripts that were present in two or more AML samples but were absent in differentiated cells (Supplementary Table 2). Most of these were short transcripts and only 28 had evidence of splicing into exons of annotated genes. RT-qPCR and/or CAGE analyses on primary samples would be required to validate such alternative TSSs emerging from A-DAR elements, especially given that only a subset are supported by GENCODE or FANTOM5 annotations (Supplementary Table 2). Nevertheless, one notable example involved a LTR2C element active in a subset of AMLs, which acted as a non-reference promoter for *SAGE1* (Figure 2B), a known cancer/testis antigen^39,40^. Another example is an LTR2B element that is active in the majority of AML samples and is an annotated promoter of the *RHEX* gene. *RHEX* regulates erythroid cell expansion^41^ and is highly expressed in AML (not shown).

We then asked whether A-DARs are marked by promoter- or enhancer-associated histone modifications. Using ChIP-seq data from the Blueprint consortium (n samples: 29 AML, 7 macrophage, 8 monocyte), we first plotted the percentage of elements from each ERV family that were marked by H3K27ac, H3K4me1, H3K4me3 or H3K9me3 in AML and differentiated myeloid cells (Supplementary Figure 3A). Notably, in AML samples an average 5.7-15.2% of elements from each family overlapped H3K4me1 peaks, a mark predominantly associated with poised and active enhancers. This was substantially higher than the fraction overlapping with the active promoter mark H3K4me3 (1.3-3.4%). Indeed, a more detailed analysis of histone modification patterns at A-DAR elements showed that H3K4me1 is either found in conjunction with H3K27ac (active enhancers), or on its own (primed enhancers) but is rarely found together with H3K4me3 (Figure 2C, Supplementary Figure 3B). Clustering analysis of these patterns demonstrated that while some elements within a family bear active marks in both AML and differentiated cells, a substantial portion (10-37%, depending on the family; median 20%) display enhancer-like profiles only in AML samples (Figure 2C, Supplementary Figure 3B). ChIP-seq profiles confirmed that these AML-specific elements had elevated H3K4me1 and H3K27ac in AML when compared to differentiated cells (Figure 2D). A total of 1,122 and 411 A-DAR elements were marked by H3K4me1 and H3K27ac respectively (333 had both marks), in at least two AML samples. A-DARs are therefore frequently associated with enhancer-like profiles in AML.

**Figure 3.**
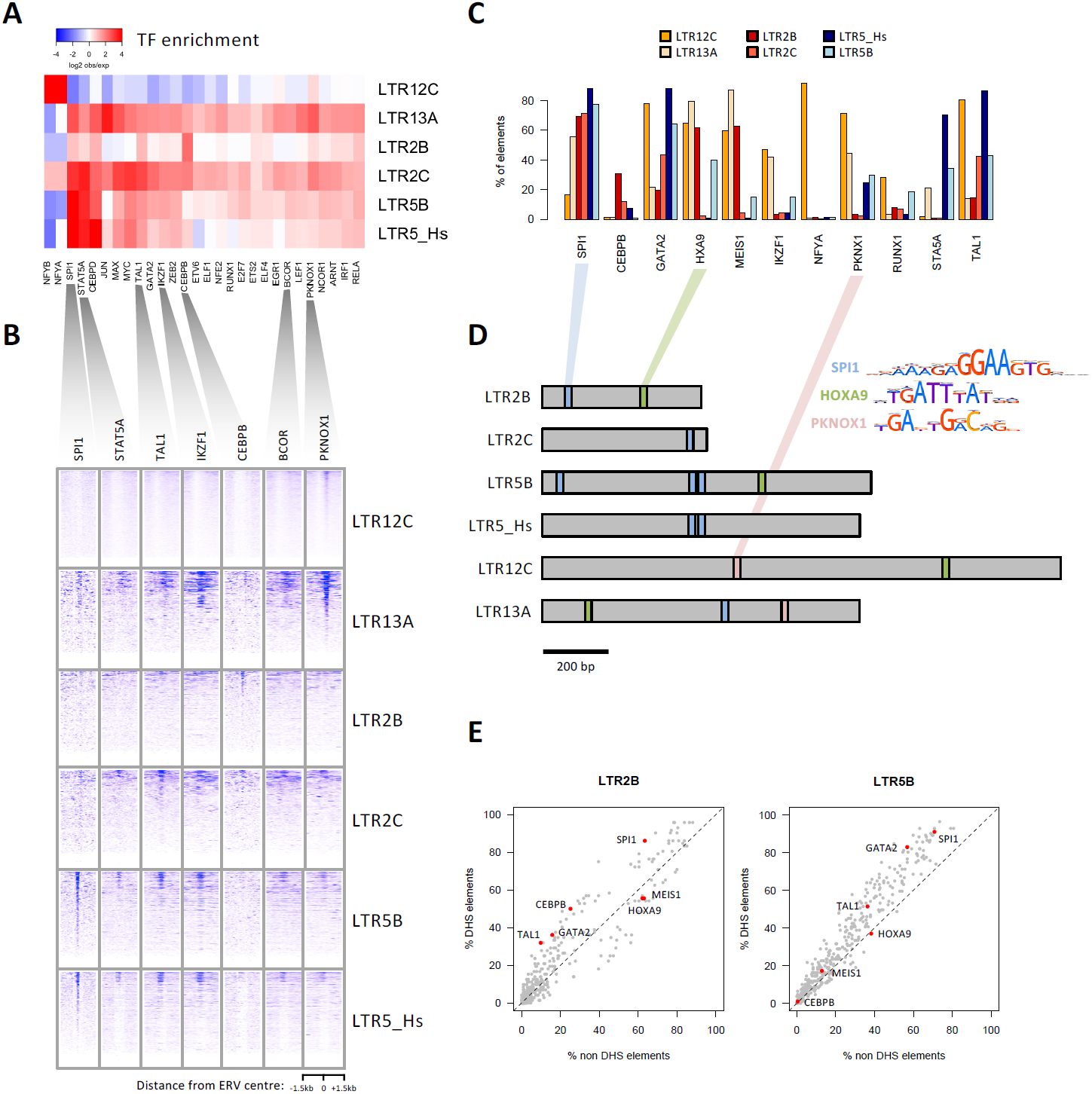
A-DARs bind AML-related transcription factors (TFs). **A.** Heatmap of the observed/expected enrichment for TF binding sites in K562 cells. **B.** ChIP-seq profiles of selected TFs across all elements of each A-DAR family. For each family, elements are displayed in the same order across all TF profiles. **C.** Percentage of ERVs from each family bearing a binding motif for the indicated TFs. **D.** Location of selected TF motifs at the consensus sequences of each A-DAR family. **E.** TF motif frequency at LTR2B and LTR5B elements, comparing those that overlap DHSs with those that do not.

To test whether myeloid leukemia cell lines could be used to dissect the putative enhancer roles of A-DARs, we performed H3K27ac ChIP-seq on HL-60, MOLM-13, OCI-AML3 and K562 cells, and compared patterns with those seen in AML samples. A-DAR elements that overlap H3K27ac peaks in AML samples were also frequently associated with this mark in cell lines (Figure 2E). A ChromHMM annotation for K562 cells from ENCODE further supported that these elements often bear enhancer signatures (Figure 2E). It is worth noting that there is substantial variation in H3K27ac enrichment of A-DARs among cell lines, much like in primary AML samples. Nonetheless, example loci show that H3K27ac deposition at A-DAR elements in cell lines can recapitulate primary AML data (Figure 2F), opening up the opportunity to functionally test for enhancer activity of these loci in cell lines.

### A-DARs bind AML-related TFs

Previous ChIP-seq or motif analyses had identified several TFs associated with the ERV families identified here^15,17^. These included hematopoiesis- and AML-related TFs such as TAL1, SPI1, GATA2 and ARNT, amongst others. To confirm and extend these observations, we first performed our own analysis of TF ChIP-seq data from K562 cells (ENCODE consortium). Our comparison with AML data above gave us confidence that K562 cells were an adequate model to study TF binding patterns at A-DARs. We analysed all TF ChIP-seq peak data available from ENCODE and selected TFs that are bound to at least 5% of the elements in a given ERV family, in a statistically significant manner, yielding a list of 217 TFs (Figure 3A; Supplementary Table 3). The vast majority of these TFs were found to be expressed in AML samples (198 had higher expression than TBP) and many of them are involved in hematopoietic gene regulation and/or in the etiology of AML, including SPI1, TAL1, IKZF1 and PKNOX1 (Figure 3A). ChIP-seq profiles of individual elements revealed a localized pattern of TF binding at a subset of elements (Figure 3B), with different ERV families binding different combinations of TFs. To evaluate TF binding in a primary cell type, we analysed data from CD34+ hematopoietic progenitors, from the BloodChIP database^42^. This revealed clear binding enrichment for FLI1, GATA2, LYL1, RUNX1 and TAL1 in at least one of the ERV families (Supplementary Figure 4).

**Figure 4.**
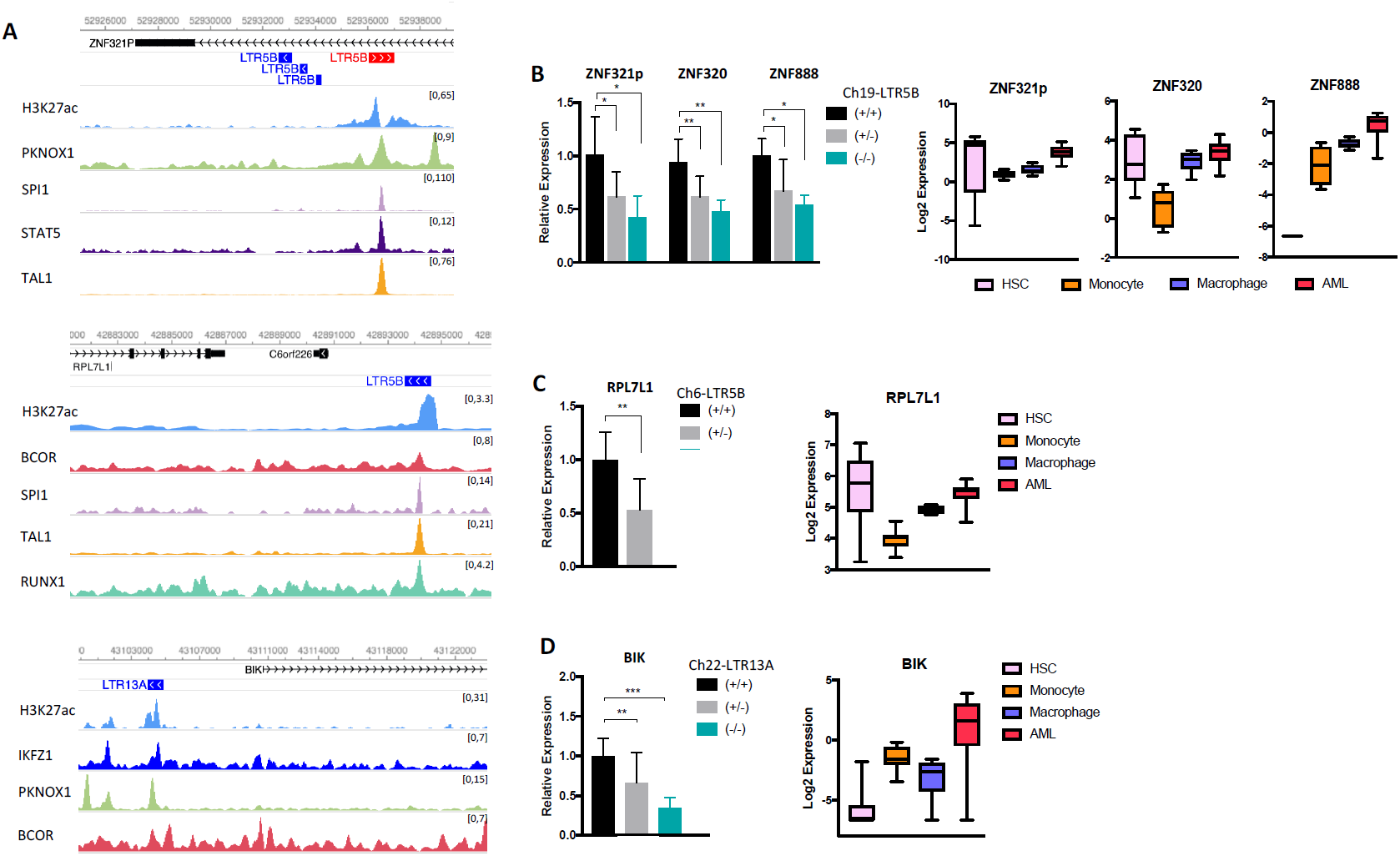
Regulatory ERVs modulate host gene expression. **A.** Genome browser view of three candidate ERVs, showing H3K27ac and TF ChIP-seq tracks in K562 cells. **B**,**C**,**D.** Expression of nearby genes (left) in the excision clones of the indicated ERVs. The error bars show standard deviation (n≥4 technical replicates for each independent clone: 2 +/+, 3 +/- (B,C,D) and 1 -/- (B,D); ANOVA with Tukey’s multiple comparison test, \**p* < 0.05, \*\**p* < 0.01, \*\*\**p* < 0.001) and (right) in HSC (n=6), monocyte (n=8), macrophage (n=6) and AML (n=27) samples (boxes indicate median ± IQR +, whiskers indicate range).

We also performed TF motif analysis (Figure 3C, Supplementary Table 4), which were largely congruent with the ChIP-seq data. Apart from confirming the presence of motifs for SPI1, PKNOX1 and other TFs, in four different ERV families we found enrichment for HOXA9/MEIS1 motifs, co-expression of which is sufficient to drive leukemogenesis in mouse models^43^. In line with the high frequency of many of the identified TF motifs, we found that they were present in the consensus sequences of each ERV family (Figure 3D), suggesting that the respective retroviruses brought in these motifs within their LTRs upon invasion of the human genome. Finally, we asked whether some TF motifs were responsible for chromatin opening at individual elements. We tested for motif enrichment in elements with DHSs (DHS+) in at least five of the analysed AML samples, when compared to DHS-negative elements (Supplementary Data 1). In four of the ERV families we identified several enriched motifs (none in LTR2C or LTR13A), such as TAL1 (in LTR2B, LTR5_Hs and LTR12C), CEBPB (in LTR2B) and GATA2 (in LTR5B and LTR12C). However, the differences in motif frequency between DHS+ and DHS-elements were modest, making TF motifs poor discriminators of these two groups (Figure 3E, Supplementary Figure 5). For example, even though SPI1 binding motif is present in the majority of DHS+ elements, a large portion of non-DHS elements also harbor this motif (Figure 3E). This suggests that other factors play a role in determining LTR regulatory potential, in line with our previous observations in mouse stem cells^44^.

**Figure 5.**
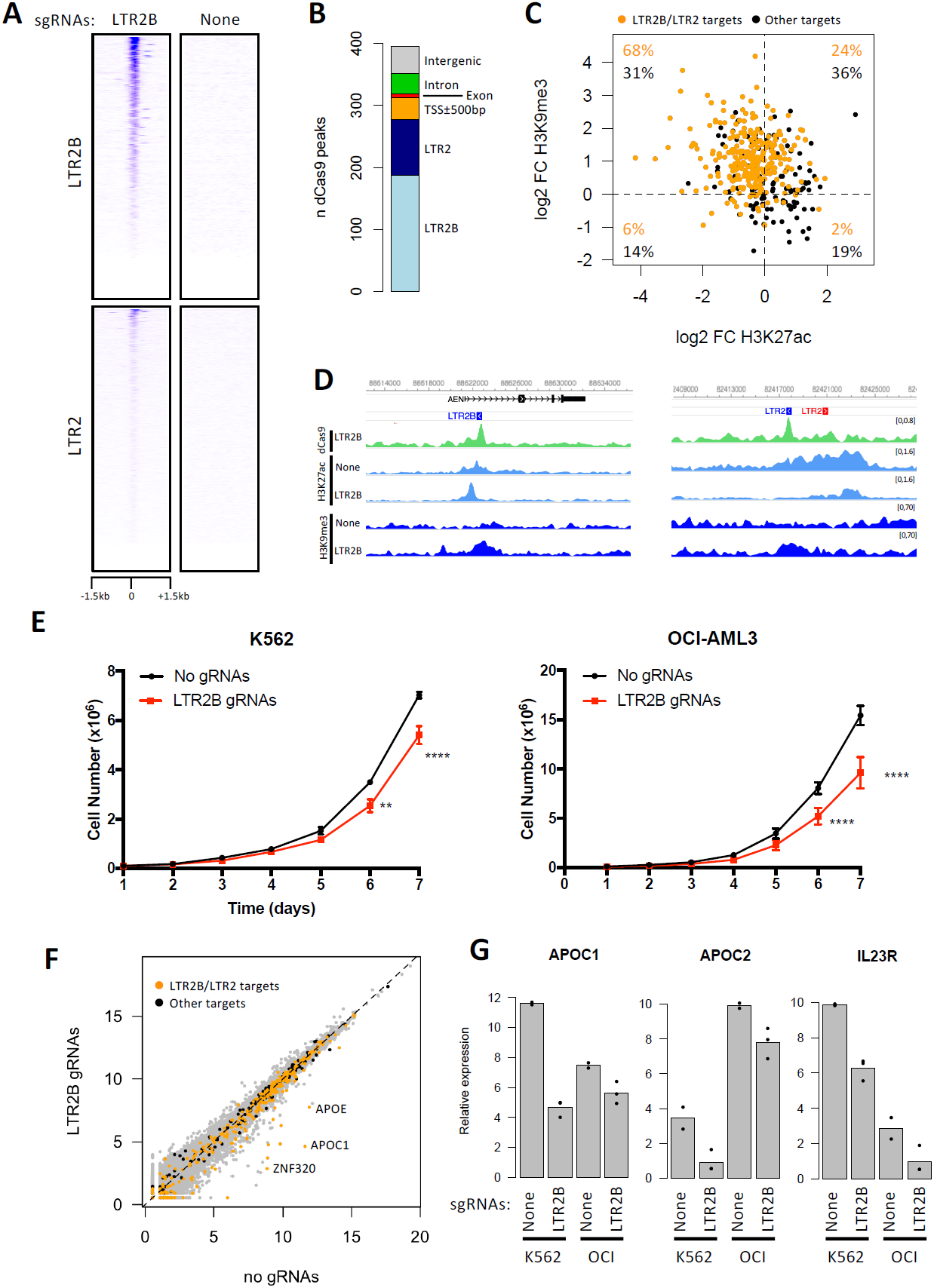
CRISPRi provides robust and selective targeting of LTR2B and LTR2 elements, which led to impaired cell growth. **A.** Profile of dCas9 ChIP-seq signal over LTR2B and LTR2 elements in K562 cells expressing LTR2B sgRNAs or an empty vector (‘None’). **B.** Number of dCas9 peaks overlapping LTR2 and LTR2B elements, or other genomic features. **C.** Log2 ratio of the ChIP-seq signal at dCas9 peaks (1kb regions from the centre of each peak) between K562 cells expressing LTR2B sgRNAs or empty vector. Orange points highlight dCas9 peaks overlapping LTR2B or LTR2 elements. **D.** Two examples of LTR2B/LTR2 elements targeted by dCas9, showing decreased H3K27ac and increased H3K9me3. **E.** Cell proliferation assay in K562 (left) and OCI-AML3 (right) cells expressing LTR2B sgRNAs or an empty vector (2 biological replicates and at least 2 technical replicates from each biological replicate were used, ANOVA with Sidak’s multiple comparison test, \*\**p* < 0.01, \*\*\*\**p* < 0.0001). **F.** Gene expression levels in K562 cells expressing LTR2B sgRNAs or empty vector. Orange points highlight genes within 50kb of a dCas9 peak targeting LTR2B/LTR2 elements; black points refer to genes within 50kb of other dCas9 peaks. **G.** Comparison of expression changes at selected genes between K562 and OCI-AML3 (‘OCI’) CRISPRi cells (n=3 biological replicates).

These analyses suggest that the potential regulatory activity at particular ERV families in AML is likely driven by the binding of hematopoiesis-associated TFs, which are either upregulated in AML or whose binding sites become accessible in AML through epigenetic alterations.

### Genetic excision of A-DAR elements interferes with host gene expression

To test for causal roles of enhancer-like A-DAR elements in gene regulation, we used CRISPR-Cas9 to delete three candidate ERVs (Supplementary Figure 6). The selected ERVs are enriched in H3K27ac, bound by multiple hematopoiesis-associated TFs in K562 cells (Figure 4A), and overlap DHSs in multiple AML samples, but not in monocytes or macrophages (Supplementary Figure 7). We generated clones with heterozygous or homozygous deletions of these ERVs in K562 cells and measured the expression of associated genes in multiple clones. Other leukemia cell lines (HL-60, OCI-AML3, MOLM-13) proved more refractory to genetic deletion, due to the low efficiency of Cas9 delivery and single cell expansion.

**Figure 6.**
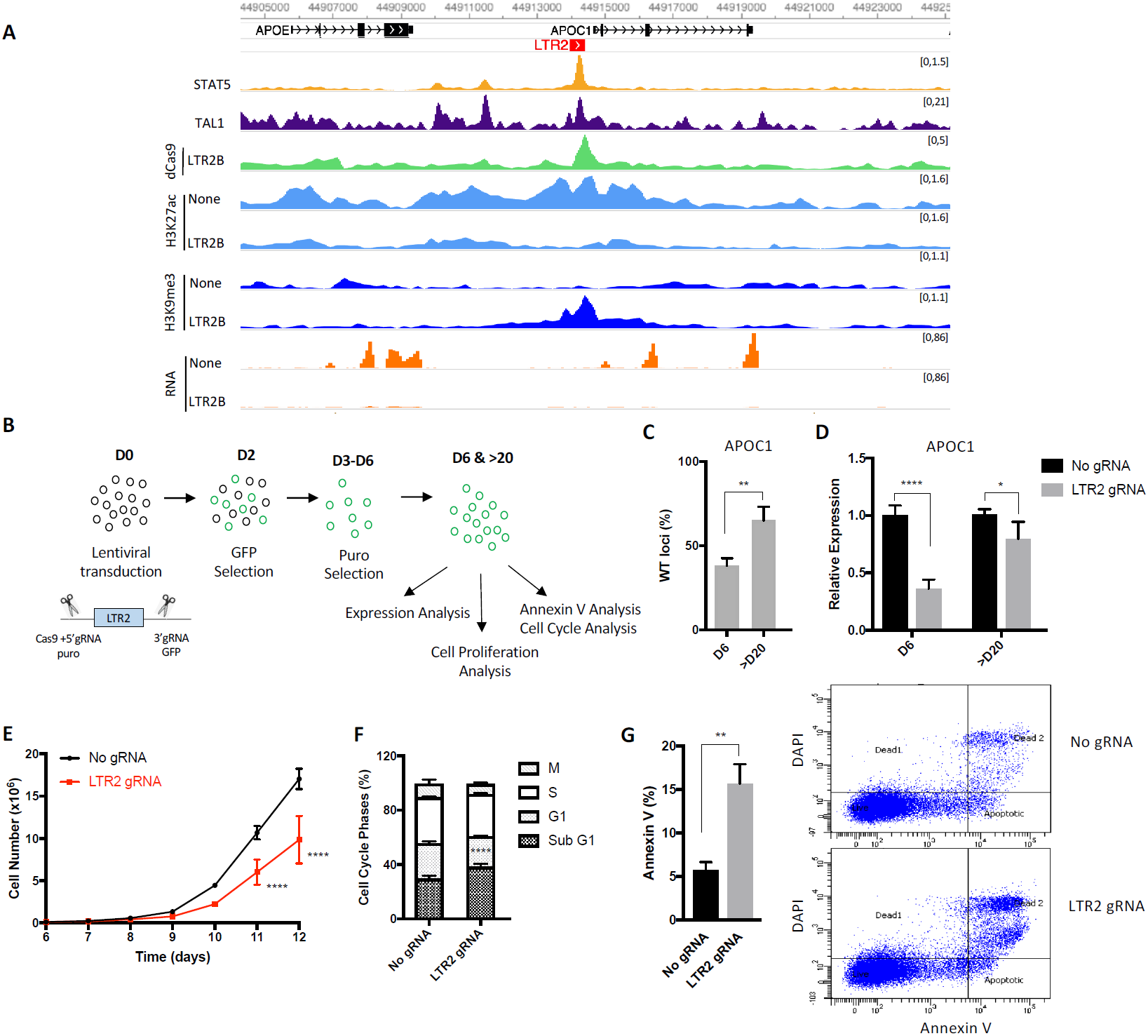
*APOC1*-LTR2 element regulates *APOC1* expression and is essential for cell proliferation. **A.** Genome browser snapshot for *APOC1*-LTR2 element, showing TAL1, STAT5 ChIP-seq tracks for WT; H3K27ac, H3K9me3 ChIP-seq and RNA-seq tracks for no control and CRISPRi K562 cells. **B.** Schematic of the experimental design to genetically excise *APOC1*-LTR2 element. **C.** qPCR and **D.** RT-qPCR data from cells with *APOC1*-LTR2 excision. The error bars show standard deviation (n≥3 biological replicates, t-test (C) ANOVA with Tukey’s multiple comparison test (D), \**p* < 0.05, \*\**p* < 0.01, \*\*\*\**p* < 0.0001). **E.** Cell proliferation assay of control and *APOC1*-LTR2 excised cells after puromycin selection (day 6) (n≥3 biological replicates, ANOVA with Tukey’s multiple comparison test, \*\*\*\**p* < 0.0001). **F.** Cell cycle profiles of control and *APOC1*-LTR2 excised cells. The error bars show standard deviation (n=3 biological replicates, ANOVA with Tukey’s multiple comparison test, \*\*\*\**p* < 0.0001). **G.** % of Annexin V stained cells in K562 cells upon *APOC1*-LTR2 excision. The error bars show standard deviation (left, n=3 biological replicates, t-test, \*\**p* < 0.01). Representative flow cytometry analysis of Annexin V (right).

One of the deleted loci was a LTR5B element located in the first intron of *ZNF321P*, which is bound by PKNOX1, SPI1, STAT5 and TAL1 (Figure 4A, top). Deletion of this element led to a significant decrease in *ZNF321P* expression and also affected the expression of two other nearby genes, *ZNF320* and *ZNF888* (Figure 4B, left). Notably, all three genes display higher expression in AML samples when compared to monocytes and macrophages (Figure 4B, right). Interestingly, *ZNF320* is also upregulated in multiple cancer types^45^. ZNF320 is a member of the Krüppel-associated box (KRAB) domain zinc finger family and predominantly binds LTR14A and LTR14B elements^46^, suggesting a potential role in ERV silencing. Heterozygous deletion of another LTR5B element, bound by BCOR, SPI1, TAL1 and RUNX1 (Figure 4A, middle), reduced the expression of Ribosomal Protein L7 Like 1 (*RPL7L1*) (Figure 4C, left), which is upregulated in AML when compared to differentiated myeloid cells (Figure 4C, right). Notably, this specific LTR5B contains a single nucleotide polymorphism (SNP) for which the minor allele (highest population frequency of 0.41) disrupts a MAFK binding motif (Supplementary Figure 7C). Using data from the GTEx project, we found that the minor allele was associated with lower *RPL7L1* expression in whole blood (Supplementary Figure 7C), suggesting that the MAFK motif is important for *RPL7L1* expression. The third deleted locus was an LTR13A element located in the vicinity of *BCL2* interacting killer (*BIK*), and is enriched for IKFZ1, PKNOX1 and BCOR binding (Figure 4A, bottom). Excision of this particular element led to around 3-fold reduction in *BIK* expression (Figure 4D, left), which is higher in AML samples when compared to other hematopoietic cell types (Figure 4D, right). This LTR13A also contains a SNP, where the minor allele (highest population frequency of 0.5) is a critical residue in a RUNX1 binding site, but that was not associated with any significant differences in *BIK* expression in whole blood (Supplementary Figure 7E). Overall, CRISPR-mediated genetic deletion assays demonstrate a direct role of individual A-DAR elements in gene regulation in K562 cells. Moreover, DHSs within the candidate ERVs and high expression of their associated genes in AML patients provide strong evidence for their regulatory activation *in vivo*.

### CRISPRi-mediated inactivation of LTR2B elements leads to growth suppression

To test the regulatory function of multiple A-DAR elements simultaneously, we next sought to epigenetically silence one ERV family by CRISPR interference (CRISPRi) using a catalytically dead Cas9 (dCas9) fused to the KRAB transcriptional repressor protein. We targeted the LTR2B family, which was the only one with AML-specific DHS enrichment and no enrichment in CD34+ cells (Figure 1B), suggesting a more cancer-specific role than other A-DARs. We designed 4 sgRNAs targeting the most conservative regions of LTR2B family, predicted to recognize around 217 copies (68%). Our LTR2B sgRNAs are also predicted to target copies of highly related LTR2 family (71 copies; 8%). To determine dCas9 specificity on a genome-wide scale, we performed dCas9 ChIP-seq in K562 cell lines expressing LTR2B sgRNAs or empty vector. We detected 395 dCas9 peaks in cells with LTR2B sgRNAs (and none in control cells), 187 of which were associated with LTR2B elements, and 90 with LTR2 elements (Figure 5A,B). The remaining 118 peaks (Figure 5B) were included in downstream analyses to evaluate putative off-target effects. We performed H3K27ac and H3K9me3 ChIP-seq in the same cells to assess the epigenetic changes imparted by CRISPRi. We quantified the ratio in histone modification levels at dCas9 peaks between cells expressing LTR2B sgRNAs and those with the empty vector control. As expected, upon CRISPRi in K562 cells we observed a reduction of H3K27ac signal and/or gain of H3K9me3 signal at most loci bound by dCas9, demonstrating effective epigenetic editing (Figure 5C,D). Notably, LTR2B/LTR2 target sites generally underwent more pronounced changes in H3K27ac and H3K9me3 levels when compared to off-target sites. Changes in histone modification levels upon CRISPRi were further confirmed by ChIP-qPCR at LTR2B elements (Supplementary Figure 8A). In OCI-AML3 cells we observed a similar trend in epigenetic alterations upon CRISPRi, albeit to a lesser extent than in K562 cells (Supplementary Figure 8B,C).

Intriguingly, proliferation assays showed that epigenetic silencing of LTR2B and LTR2 elements by CRISPRi significantly suppressed cell proliferation in both K562 and OCI-AML3 cell lines (Figure 5E). To test the impact of LTR2B and LTR2 inactivation on the host transcriptome, and gain insights into the mechanism underlying impaired cell growth, we performed RNA-seq in both cell lines (Figure 5F; Supplementary Figure 8D). We identified a total of 58 and 99 differentially expressed genes in K562 and OCI-AML3 cells, respectively (Supplementary Table 5). To elucidate direct effects of CRISPRi, we focused on genes that are within 50 kb of a dCas9 peak and found 15 and 6 differentially expressed genes (in K562 and OCI-AML3 cells, respectively), all but one of which were downregulated. Only one of these genes (*BIK*), which was downregulated in OCI-AML3, was associated with an off-target dCas9 peak. The remaining genes were associated with 15 different LTR2B/LTR2 elements. Four of these elements were intronic and thus we cannot exclude the possibility that dCas9 binding interfered with transcriptional elongation^47^.

In some instances, the LTR2B/LTR2 element was very close to the promoter of the affected gene, such that silencing could have resulted from H3K9me3 spreading. We therefore performed genetic deletion of one of these elements, which also led to a decrease in expression of the adjacent *ZNF611* gene, albeit to a lesser extent than by CRISPRi (Supplementary Figure 8E). Several genes displayed decreased expression in both cell lines (Figure 5G), although only apolipoprotein C1 (*APOC1*) reached statistical significance in both contexts. Notably, five apolipoprotein genes were downregulated in at least one of the cell lines. *APOC1, APOC2, APOC4-APOC2* and *APOE* lie within a cluster on chromosome 19, and may all be controlled by the same LTR2 element, located upstream of *APOC1*. On the other hand, *APOL1* is on chromosome 22 and close to an LTR2B insertion. Given the key roles that lipid metabolism plays in supporting cancer cell survival^48^, the coordinated downregulation of apolipoprotein genes could underpin the reduced cell growth observed upon silencing of LTR2B/LTR2 elements in leukemia cell lines.

Overall, these data show that a subset of LTR2B and LTR2 elements act as key gene regulators in leukemia cell lines, and that their epigenetic silencing impairs cell growth, providing evidence for a functional role in AML.

### *APOC1*-associated LTR2 is required for proliferation of myeloid leukemia cells

APOC1 has recently been shown to maintain cell survival in AML and the knockdown of *APOC1* impairs cell growth^49^. Similar findings were made in pancreatic and colorectal cancer, where *APOC1* overexpression is associated with poor prognosis^50,51^. We therefore asked whether ERV-mediated regulation of *APOC1* could affect cell growth. There is an LTR2 insertion upstream of the *APOC1* promoter (*APOC1*-LTR2; Figure 6A, Supplementary Figure 9A,B), which has been previously described to act as an alternative promoter in several tissues, but only accounts for up to 15% of total *APOC1* transcription^52^. In K562 and OCI-AML3 RNA-seq data, we found no evidence of *APOC1*-LTR2 promoter activity (Figure 6A, Supplementary Figure 8A), which we have confirmed by RT-qPCR (Supplementary Figure 9C), suggesting that *APOC1*-LTR2 could act as an enhancer element. *APOC1*-LTR2 is enriched in STAT5 and TAL1 binding and shows an increase in H3K9me3 and decrease in H3K27ac upon CRISPRi in both K562 and OCI-AML3 (Figure 6A, Supplementary Figure 9B).

To test for a direct role of *APOC1*-LTR2 in *APOC1* gene expression and cell growth, we deleted this element in K562 cells without affecting the *APOC1* promoter (Supplementary Figure 9A). We obtained 7 heterozygous and 8 homozygous clones from a total of 110 clones. Interestingly, none of the homozygous clones were able to grow more than 10 days in culture, suggesting that homozygous deletion may impair cell growth. To pursue the impact of *APOC1*-LTR2 on cell growth, we used lentiviral-mediated CRISPR-Cas9 delivery and performed assays in a pool of edited cells (Figure 6B). At day 6, after GFP and puromycin selection of the two flanking sgRNAs, we observed around 60% deletion of *APOC1*-LTR2 and more than 2.5-fold reduction in *APOC1* gene expression compared to an empty vector control (Figure 6C,D). Deletion of *APOC1*-LTR2 also led to decrease in the expression of the nearby *APOE* gene (Supplementary Figure 9D), consistent with the results from CRISPRi (Figure 6A). Remarkably, deletion of this element was sufficient to drive a significant suppression of cell proliferation compared to control cells (Figure 6E). This is particularly notable given the partial nature of the deletion, emphasizing the dramatic growth arrest seen in homozygous null CRISPR clones. As there is a fraction of unedited cells in the pool, we asked whether the unedited cells may outcompete edited cells over time. After day 20, the deletion was reduced to around 35% and only 1.2-fold difference was observed in *APOC1* expression, and consequently there was no difference in cell proliferation, indicating that *APOC1*-LTR2 provides cells with a selective growth advantage (Figure 6C,D; Supplementary Figure 9E). To further investigate how *APOC1*-LTR2 deletion leads to impaired cell growth, we analyzed cell cycle and apoptosis with flow cytometry in K562 cells at day 6. While no differences in G1, S, and G2 phases were detected, there was a significant increase in the sub-G1 population in edited cells (Figure 6F). In agreement with this, Annexin V signal was significantly higher in edited cells compared to unedited cells at day 6 (Figure 6G), showing that the deletion of *APOC1*-LTR2 induces apoptosis, which is in line with known effects of *APOC1* depletion^49-51^. As expected, this difference is much smaller after day 20 (Supplementary Figure 9F). We also tested the effect of *APOC1*-LTR2 deletion in OCI-AML3 cells, but due to the low efficiency of Cas9 delivery and low viability of cells at day 6, we performed expression and annexin V analysis at day 10.

Similar to what we observed in K562 cells, *APOC1*-LTR2 deletion in OCI-AML3 cells led to around 4-fold decrease in *APOC1* expression and increased annexin V signal, and these effects were milder at day 23 (Supplementary Figure 9G,H). Our findings indicate that the *APOC1*-LTR2 element is essential for proliferation of leukemia cells by acting as an enhancer of the *APOC1* gene, which in turn controls cell survival via an anti-apoptotic mechanism.

Notably, DNase-seq peaks associated with *APOC1*-LTR2 in AML samples are subtler than those observed in cell lines, yet a few AML samples express relatively high levels of *APOC1* (Supplementary Figure 10A,B). Interestingly, overall survival curves based on TCGA data suggest that a small proportion of patients with high *APOC1* expression have a poorer prognosis, a pattern that is also seen in patients with high APOE expression (Supplementary Figure 10C,D).

## Discussion

Here we demonstrate that particular ERVs are used as regulatory elements to activate gene expression in AML, which may be exploited by cancer cells to help drive disease phenotypes and cancer progression. Many of these ERVs are also active in CD34+ progenitor cells and are therefore not cancer-specific, but they may nonetheless be used to support a gene expression programme that blocks cellular differentiation, a key hallmark of AML. Genetic and epigenetic perturbation experiments such as the ones presented here, allow us to distinguish between ERVs that support oncogenesis and those whose activation is secondary to cellular de-differentiation.

It had been previously postulated that the epigenetically relaxed state of cancer cells provides a window of opportunity for ERV activation, triggering their intrinsic regulatory capacity^9,24,53^. However, to the best of our knowledge, all examples to date supporting this hypothesis have involved activation of cryptic promoters to drive expression of adjacent genes^24,27^. Whilst we uncovered some examples of chimeric transcripts starting from ERVs in AML (e.g., LTR2C-*SAGE1*, LTR2B-*RHEX*), which are not present in differentiated myeloid cells, our analyses suggest that active A-DARs mainly harbour chromatin signatures of enhancers.

We identified multiple ERV elements with strong evidence supporting their role as bona fide gene regulators: 1) we found striking correlations between differential chromatin accessibility at 20 ERVs and the expression of nearby genes, some of which have been linked to AML prognosis (Figure 1E,F); 2) CRISPR-mediated genetic editing experiments revealed an additional five ERVs that act as enhancers in leukemia cells (Figure 4, Supplementary Figure 8E, Figure 6); 3) CRISPRi identified another 13 different elements whose epigenetic silencing led to the down-regulation of nearby genes (Supplementary Table 6). A more exhaustive search would likely have revealed additional regulatory elements, namely via epigenetic silencing of other ERV families. Moreover, given the heterogeneity of the disease, inclusion of additional primary AML data or a focus on specific AML subtypes may have uncovered other ERV families/loci of interest.

Despite the growing evidence that ERVs can act as regulatory elements in different cancers, there are limited examples for their inappropriate activation contributing to oncogenesis, a term coined as onco-exaptation^54^. The term has been frequently used to describe the gain of regulatory activity at TEs in cancer. Our view is that, similar to the term exaptation^55^, onco-exaptation requires that this new regulatory activity provides the cancer cell with a selective advantage. Strong demonstrations of such adaptive roles are scarce. Notably, the Wang lab recently showed that an AluJb element acts as an oncogenic promoter to drive *LIN28B* expression and tumour progression in lung cancer^27^. In our study we identified an LTR2 element, the genetic and epigenetic perturbation of which suppressed cell growth and induced apoptosis of leukemia cell lines by altering lipid-related *APOC1* expression. Despite the striking cellular phenotype in cell lines, comprehensive analyses of primary AML samples are warranted to demonstrate whether these regulatory ERVs are sufficient to provide survival advantages for cancer cells *in vivo* and contribute to prognosis of AML.

Notably, we observed that AML patients with high APOC1 or APOE expression demonstrate significantly lower overall survival rate. A considerably larger number of patients would be necessary to confirm this finding, although independent datasets have led to similar observations in colorectal and pancreatic cancer^50,51^. APOC1 is also activated in monocyte-to-macrophage differentiation^56^, raising the possibility that *APOC1*-LTR2 may play other roles in haematopoiesis outside of AML.

Given their repetitive nature, one intriguing question is why particular ERVs within a family are recurrently activated in AML to drive nearby gene expression, yet the majority of them are functionally neutral. One explanation lies in the nature of inter- and intra-cellular epigenetic heterogeneity that increases during malignancy formation. This gives rise to epigenetic activation of a set of ERVs, as proposed in the epigenetic evolution model^24^. Accordingly, cells harbouring activated ERVs that drive oncogenes gain a selective advantage and increase in frequency during cancer evolution. Therefore, clonal expansion of these cells will enable the detection of oncogenic ERVs in a cell population. However, whether ERV activation contributes to cancer evolution or is simply a consequence of the molecular state of cancer remains a matter of debate.

Irrespective of whether epigenetic heterogeneity at ERVs contributes to tumour evolution, distinct patterns of ERV activity are observed across different AML patients (Supplementary Figure 1A). These differences appear to be partly driven by the underlying mutational profiles. We also identified a SNP within an ERV that seemingly affects its regulatory activity by altering a TF binding site (Supplementary Figure 7C), suggesting that genetic variation within ERVs also contributes to inter-individual differences in ERV activity. Finally, younger ERVs such as LTR5_Hs are structurally polymorphic within the human population^12,57^, adding another layer of genetic variation. Regulatory ERVs may therefore foster genetic, epigenetic and transcriptional heterogeneity of the disease with potential to contribute to clinical outcomes. One significant consequence of the molecular heterogeneity of AML is the escape of resistant clones from treatment, resulting in high relapse rates. It will be therefore interesting to discover to which extent the ERV-derived heterogeneity contributes to inter-individual differences in response to AML therapies.

Our work reveals the first example of ERVs as potentially oncogenic enhancers in AML or in any human malignancy. These data highlight the significance of expanding the search for oncogene drivers to the repetitive part of the genome, which may pave the way for the development of novel prognostic and therapeutic approaches.

## Methods

### Cell culture and cell proliferation assays

293T cells and human leukemia cell lines K562, OCI-AML3, MOLM-13 and HL-60 were routinely cultured in RPMI 1640 (and DMEM (HEK293T) supplemented with 10% fetal bovine serum, 2mM glutamax and 1% penicillin/streptomycin at 37°C in 5% carbon dioxide. Cells were maintained and split every 2-3 days.

For cell proliferation assays, exponentially growing cells were plated in 24-well plates (1×10^5^ cells/ml). Every 2-3 days media were replaced and cells were split to 1×10^5^ cells/ml. The viable cells were counted daily for 6 days.

### Cell-cycle and apoptosis assays

Cell cycle assay was performed using muse cell cycle kit by following the manufacturer’s instructions (Millipore) and the cells were analysed by BD FACS Canto II. For apoptosis assay, the cells were stained by an annexin V 647 (Thermofisher Scientific) and DAPI and analysed by BD FACS Canto II.

### CRISPR-Cas9-mediated LTR disruption

For CRISPR/Cas9 deletion of LTRs, sgRNA oligonucleotides (Sigma-Aldrich) targeting upstream and downstream of LTRs of interest were annealed and cloned into modified eSpCas9 (1.1) vector (Addgene 71814, deposited by Feng Zhang), which expresses GFP. K562 cells were nucleofected with eSpCas9 plasmid containing gRNAs using amaxa nucleofector kit V. 2 days later, cells expressing GFP were sorted on a FACS Aria II and single cells were plated onto a 96-well plate. After 2 weeks, cells were genotyped by PCR and the gene expression of LTR-knockout cells was analyzed by RT-qPCR.

For LTR2-APOC1 deletion, 5’ sgRNAs (Sigma-Aldrich) were cloned into lentiCRISPR v2 (Addgene 52961) and 3’ sgRNAs were cloned into lenti_sgRNA_EFS_GFP (Addgene 65656) vector. OCI-AML3 and K562 cells were transduced with the lentiviral vectors containing sgRNAs and selected for GFP and puro. % of WT loci was determined by qPCR using APOC_R and APOC_I genotyping primers listed in Supplementary Table 7. The cells were cultured around three weeks for RNA expression and phenotypical analysis.

### CRISPRi-mediated silencing of LTRs

sgRNAs (Sigma-Aldrich) targeting multiple LTR copies were cloned into lentiviral expression vector pKLV-U6gRNA(BbsI)-PGKpuro2ABFP (Addgene 50946, deposited by K. Yusa). For LTR silencing, OCI-AML3 and K562 cells were first transduced with the lentiviral vector pHR-SFFV-KRAB-dCas9-P2A-mCherry (Addgene 60954, deposited by Jonathan Weissman), sorted for mCherry on a FACSAria II. Cells expressing mCherry were then subsequently transduced with the lentiviral sgRNA expression vector. 2 days later, the cells expressing both mCherry and BFP were sorted and cultured for transcriptional and chromatin analyses.

### Lentiviral production and transduction

Lentivirus was produced in HEK 293 T cells by triple transfection with delivery vector and the packaging plasmids psPAX2 and pMD.G. The viral supernatants were collected 48 h after transfection and filtered through a 0.45 μM filter. Target cells were transduced with lentiviral supernatant supplemented with 4 µg/mL polybrene.

### RNA Isolation and RT-qPCR

RNA was extracted using AllPrep DNA/RNA mini kit (Qiagen 80204) and DNAse treated with the TURBO DNA-free(tm) Kit (Ambion, AM1907). RNA (1 µg) was retrotranscribed using Revertaid Reverse Transcriptase (Thermo Scientific EP0441) and the cDNA was diluted 1/10 for qPCRs using MESA BLUE MasterMix (Eurogenentec, 10-SY2X-03+NRWOUB) on a LightCycler® 480 Instrument II (Roche). A list of primers used can be found in Supplementary Table 7.

### RNA-seq Library Preparation

Ribosomal RNA-depleted RNA-seq libraries were prepared from 200-500 ng of total RNA using the low input ScriptSeq Complete Gold Kit (Epicentre). Libraries were sequenced on an Illumina NextSeq 500 with single-end 75 bp reads.

### Chromatin Immunoprecipitation

Approximately 10^7^ cells were fixed with 1% formaldehyde for 12 minutes in PBS and quenched with glycine. Chromatin was sonicated using a Bioruptor Pico (Diagenode), to an average size of 200-700 bp. Immunoprecipitation was performed using 75 µg of chromatin and 5 µg of Cas9 antibody (Diagenode #C15200229-100) or 15 µg of chromatin and 2.5 µg of H3K27ac and H3K9me3 antibody (Active Motif #3913, Diagenode #C15410193). Final DNA purification was performed using the GeneJET PCR Purification Kit (Thermo Scientific. #K0701) and DNA was eluted in 80 µL of elution buffer. This was diluted 1/10 and analysed by qPCR, using the KAPA SYBR® FAST Roche LightCycler® 480 2X qPCR Master Mix (Kapa Biosistems, Cat. KK4611). A list of primers used can be found in Supplementary Table 7.

### Library preparation and sequencing for ChIP-seq and DNase-seq

ChIP-seq and DNase-seq libraries were prepared from 1-5 ng ChIP DNA or DNase DNA samples using NEBNext Ultra II DNA library Prep Kit (Illumina). Libraries were sequenced on an Illumina NextSeq 500 with single-end or paired-end 75 bp reads.

### Chromatin accessibility assay

To asses chromatin accessibility, DNase I digestion was performed as previously described^58^. 5 million cells were resuspended in RSB buffer (10 mM NaCl, 3mM MgCl2, 10mM Tris-Cl, pH 7.4). After cell lysis, the nuclei were digested with DNase I with 0, 0.1, 2, 5, 15 and 30 U for 10 min at 37°C. Digests were inactivated by the addition of 50 mM EDTA. RNA and proteins were digestion by RNase A (0.5 mg/ml) for 15 min at at 37°C and then by proteinase K (0.5 mg/ml) for 1 h at 65 °C. DNA was purified by phenol-chloroform extraction and ethanol precipitation. The resuspended DNA was analysed by qPCR, using the KAPA SYBR® FAST Roche LightCycler® 480 2X qPCR Master Mix (Kapa Biosistems, Cat. KK4611) and chromatin digested with 15 U was selected for library preparation and sequencing.

### Primary processing of high-throughput sequencing data

Reads from high-throughput sequencing data generated here or from external datasets (Supplementary Table 8) were trimmed using first trimmed using Trim Galore. ChIP-seq and DNase-seq data were aligned to the hg38 genome assembly using Bowtie2 v2.1.0^59^, followed by filtering of uniquely mapped reads with a custom script. ChIP-seq peak detection was performed using MACS2 v2.1.1^60^ with -q 0.05; for histone marks the option --broad was used. DNase-seq peak detection was performed using F-seq v1.84^61^ with options -f 0 -t 6. RNA-seq data were mapped using Hisat2 v2.0.5^62^ with option --no-softclip. Raw read counts for each gene were generated in Seqmonk with the RNA-seq quantitation pipeline, and normalised gene expression values calculated with the variance stabilizing transformation in DESeq2^63^. BigWig tracks were generated using the bamCoverage function of deepTools2.0, with CPM normalisation and 200 bp bin size. Other processed data from Blueprint, ENCODE and other sources (Supplementary Table 8) were downloaded as peak annotations or expression values (e.g., FPKM).

### DHS enrichment at repeat families

DHSs (i.e., DNase-seq peaks) were intersected with the Repeatmasker annotation and the number of overlapped DHSs per repeat family calculated. For comparison, 1000 random controls were generated by shuffling the DHSs in a given sample, avoiding unmappable regions of the genome. P values were calculated based on the number of random controls for which the number of DHS overlaps displayed more extreme values (at either tail of the distribution) than those seen with the real DHSs. Enrichment values were calculated by dividing the number of real DHS overlaps with the mean number of DHS overlaps in the random controls. Significantly enriched repeat families had: 1) p<0.05, 2) >2-fold enrichment, 3) >20 copies overlapped by DHSs. Selected families were significantly enriched for DHSs in at least one of the cell lines analysed (HL-60, OCI-AML3, MOLM-13) and in >10% of AML samples.

### Mutational profile analysis

A-DAR elements overlapping DHSs in at least one sample were selected, and a correlation matrix built based on the patterns of DHS overlap between samples. These were compared with the AML mutational profiles extracted from the respective publications^5,6^. Correlation coefficients between AML samples sharing a particular mutation were compared with correlation coefficients between samples without the mutation.

### Identification of active A-DAR promoters

Aligned BAM files from Blueprint RNA-seq data were processed using StringTie v1.3.3b^64^ with options --rf -G to generate sample-specific transcriptome assemblies guided by the GENCODE annotation v26. Spliced transcripts initiating at A-DAR elements were then identified by intersecting the TSSs of multi-exon transcripts A-DAR annotations. A-DAR elements with TSSs in AML samples but not in differentiated cells were selected and the associated transcripts visually inspected to identify those with evidence of splicing into GENCODE-annotated genes. TSSs were also checked against the FANTOM5 robust CAGE peak set (hg38 version, with fairly remapped and newly identified peaks).

### K562 TF ChIP-seq analysis

ENCODE TF ChIP-seq peak files from K562 (Supplementary Table 8) were downloaded and intersected with A-DAR annotations, as well as with a randomly shuffled version of these elements. TFs significantly enriched (corrected p<0.05) in at least one of the A-DAR families, covering at least 5% of the elements in that family were selected. For each TF, average enrichment values were calculated across technical and biological replicates, as well as independent ChIP-seq experiments of the same TF.

### TF motif analysis

Motif analysis of A-DARs was performed using the AME and FIMO tools of the MEME SUITE v5.0.1^65^ using the HOCOMOCO v11 human TF motif database. Motifs enriched in at least one A-DAR family were identified using AME, and motif frequency and location extracted using FIMO. Consensus sequences were downloaded from Dfam^31^.

### CRISPRi ChIP-seq and RNA-seq analyses

Normalised H3K27ac and H3K9me3 ChIP-seq read counts were extracted around dCas9 peaks (±500 bp from the peak centre). Genes within 50 kb of a dCas9 peak were considered as putative direct targets of CRISPRi. Differential gene expression analysis was performed using DEseq2^63^.

## Acknowledgements

We thank Yasmine Benbrahim for ideas informing bioinformatic analyses, the Dawson lab for their guidance in lentiviral transduction of AML cell lines, Brian Huntly for providing OCI-AML3, MOLM-13 and HL-60 cell lines, Diego Villar and Jenny Frost for critical reading of the manuscript. This work was supported by funding from Barts Charity (Small Project Grants – MGU0462). O.D. received funding from the People Programme (Marie Curie Actions) of the European Union’s Seventh Framework Programme (FP7/2007-2013) under REA grant agreement n° 608765. M.R.B. was supported by a Sir Henry Dale Fellowship (101225/Z/13/Z), jointly funded by the Wellcome Trust and the Royal Society. This study makes use of data generated by the Blueprint Consortium. A full list of the investigators who contributed to the generation of the data is available from www.blueprint-epigenome.eu. Funding for the project was provided by the European Union’s Seventh Framework Programme (FP7/2007-2013) under grant agreement no 282510 – BLUEPRINT. This research utilised Queen Mary’s Apocrita HPC facility, supported by QMUL Research-IT^66^.

## Author Contributions

O.D. and M.R.B. designed the study and experiments and wrote the manuscript. O.D. performed cell culture, DNase-seq, ChIP-seq, RNA-seq, CRISPR, CRISPRi and cellular phenotyping. M.A. generated the ZNF611-LTR2B KO. C.D.T. assisted in the design and execution of CRISPR experiments. A.R.M. performed overall survival analyses. M.A.D. assisted in the establishment of CRISPRi cell lines. M.R.B. performed bioinformatic analyses.

## Availability of data and code

Datasets are available through the Gene Expression Omnibus (GEO) under accession number GSE136764. Scripts used for data analysis are available at https://github.com/MBrancoLab/Deniz_2019_AML.

## Supplementary Information

**Supplementary Figures 1-10**

**Supplementary Table 1:** The results of DHS permutation test for each transposon in AML cell lines, samples and other hematopoietic cell types

**Supplementary Table 2:** List of AML-associated spliced transcripts emanated from A-DARs

**Supplementary Table 3:** K562-enriched TFs that are bound to A-DARs

**Supplementary Table 4:** TF motif frequency within A-DARs with and without DHSs

**Supplementary Table 5:** Differentially expressed genes in K562 and OCI-AML3 upon CRISPRi

**Supplementary Table 6:** List of ERV elements that display regulatory activity in myeloid leukemia

**Supplementary Table 7:** Primers used in this study

**Supplementary Table 8:** External datasets used in this study

**Supplementary Data 1:** Results from the analysis of motif enrichment (MEME suite)

