## Supplementary Figures for "Endogenous retroviruses are a source of enhancers with oncogenic potential in acute myeloid leukaemia"

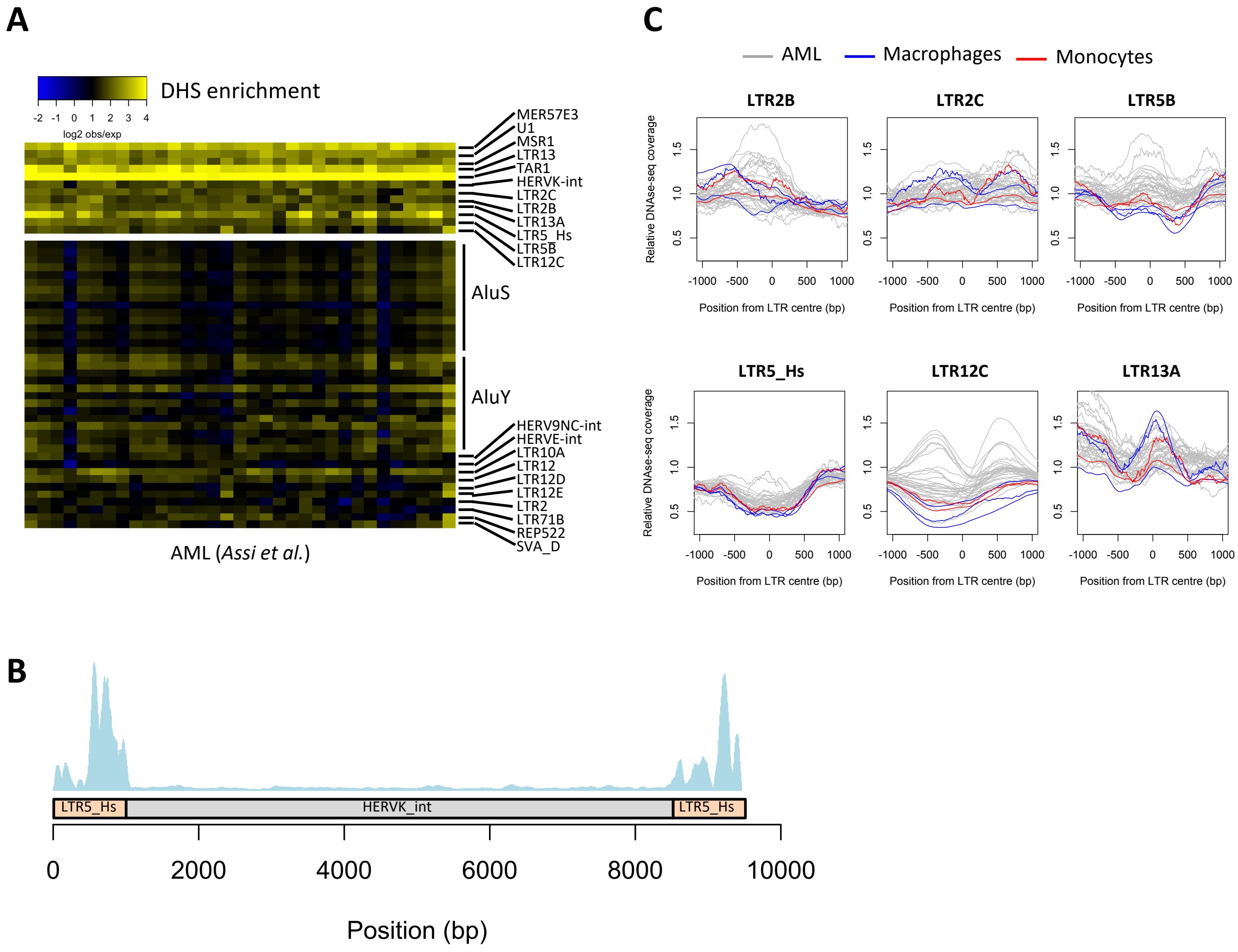


**Supplementary Figure 1. A.** Heatmap of the observed/expected enrichment for DHSs in selected repeat families, for AML sample data from Assi et al, 2018. **B.** DNase-seq profile across a representative full-length HERV-K element in OCI-AML3 cells. All DNase-seq data were aligned to this sequence, including non-unique reads. **C.** Average DNase-seq profiles for each A-DAR family across different AML, macrophage or monocyte samples.

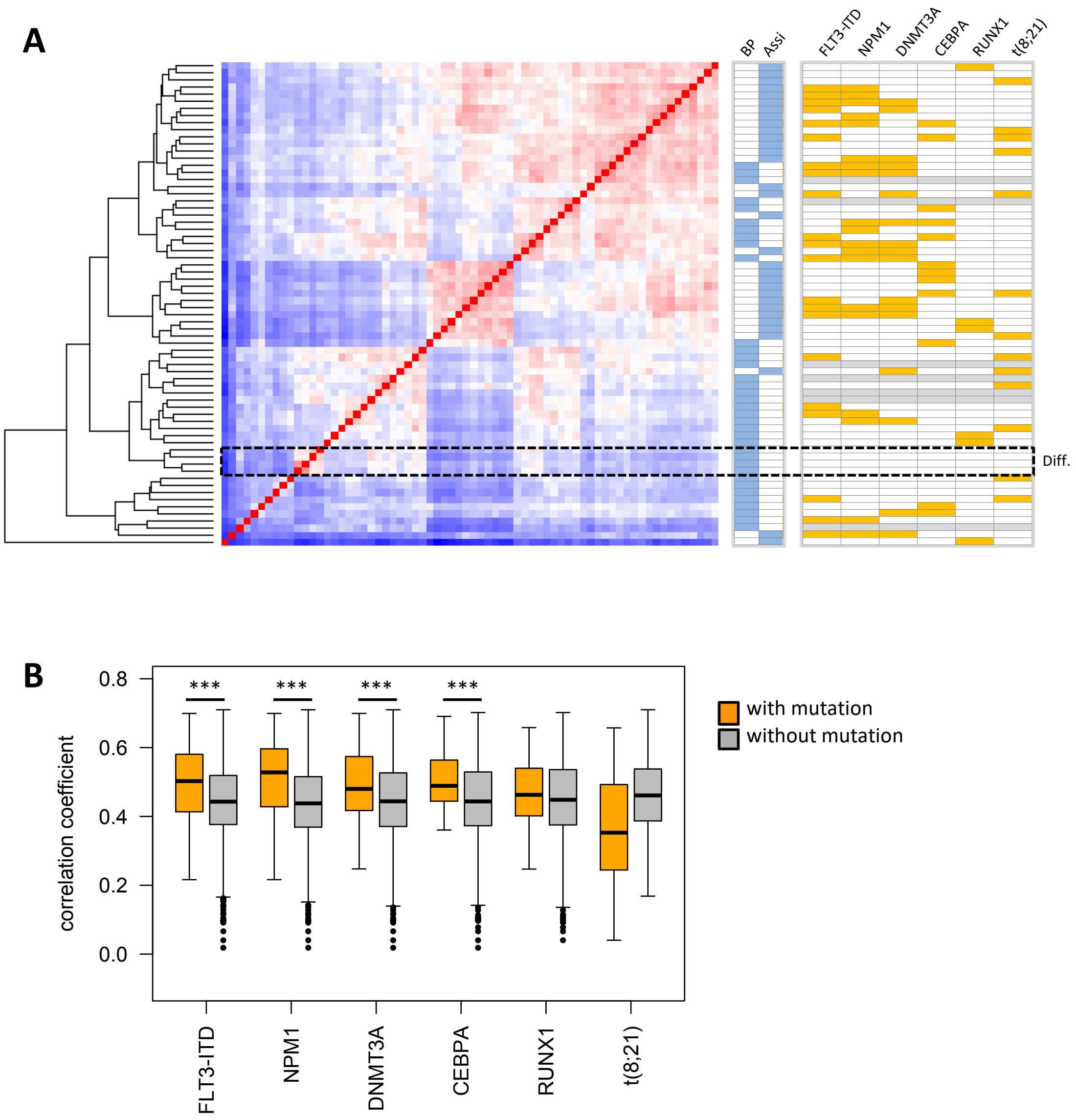


**Supplementary Figure 2. A.** Heatmap of correlation coefficients related to the presence or absence of DHSs at A-DAR elements (left). Only elements where at least one sample contained a DHS at that site were used. Genotypes and data source of each sample are shown on the right. Data were either from the Blueprint project ('BP') or from Assi et al, who also generated the genotyping data represented here. Grey lines indicate samples with no genotyping data. 'Diff.' refers to differentiated myeloid cells, i.e., macrophages and monocytes. **B.** Distribution of correlation coefficients between AML samples with a given mutation and those without (*** $p < 1E\text{-}5$, t-tests with Benjamini-Hochberg correction).

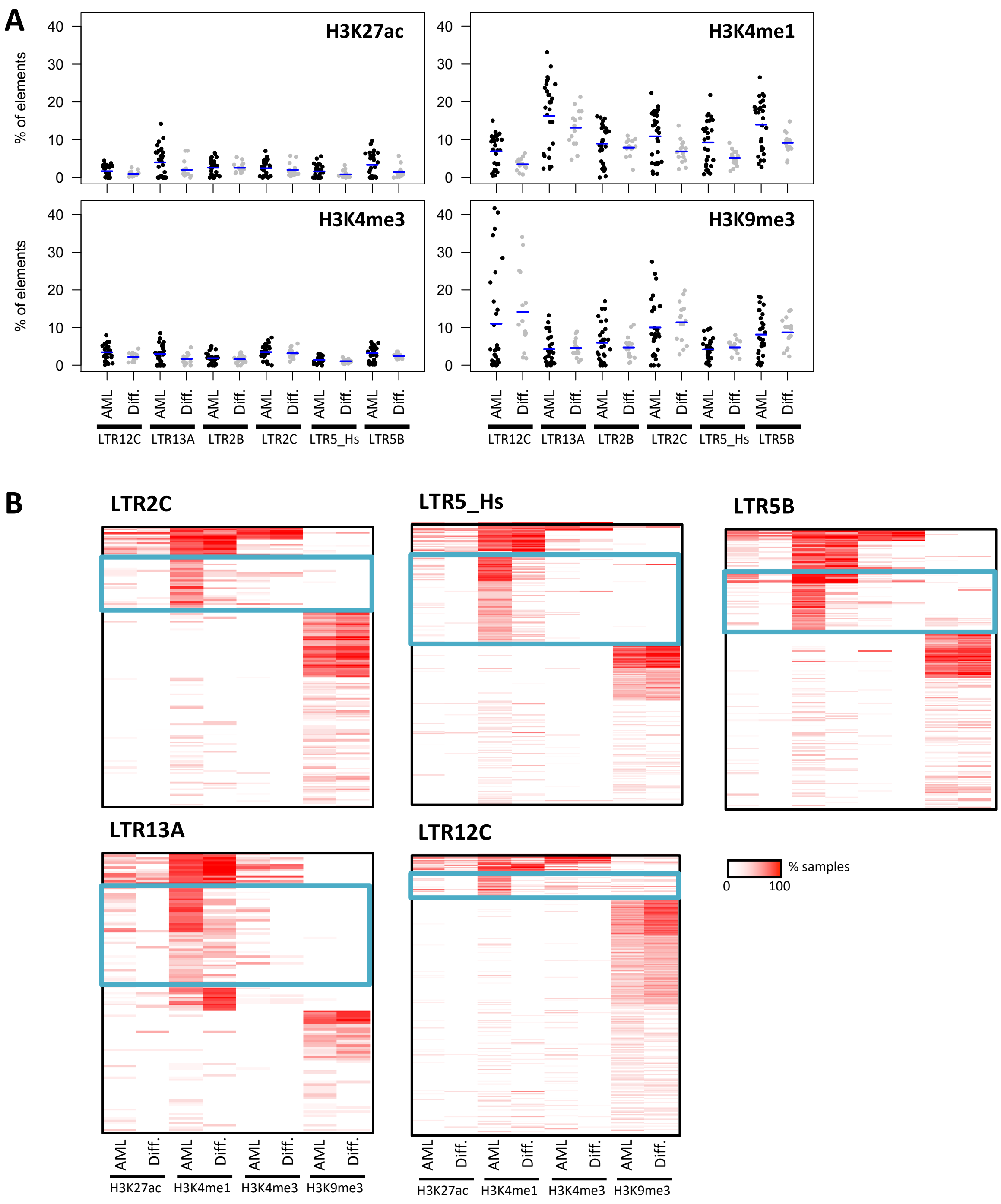


**Supplementary Figure 3. A.** Percentage of elements from each A-DAR family overlapping peaks for the indicated histone modifications. Each data point represents a different sample. **B.** Heatmap of overlap between LTR2C, LTR5B, LTR5_Hs, LTR13A or LTR12C elements and histone modification peaks. Colour intensity represents the percentage of AML or differentiated cell samples where overlap is observed. Blue boxes highlight clusters of elements associated with preferential H3K4me1 marking in AML.

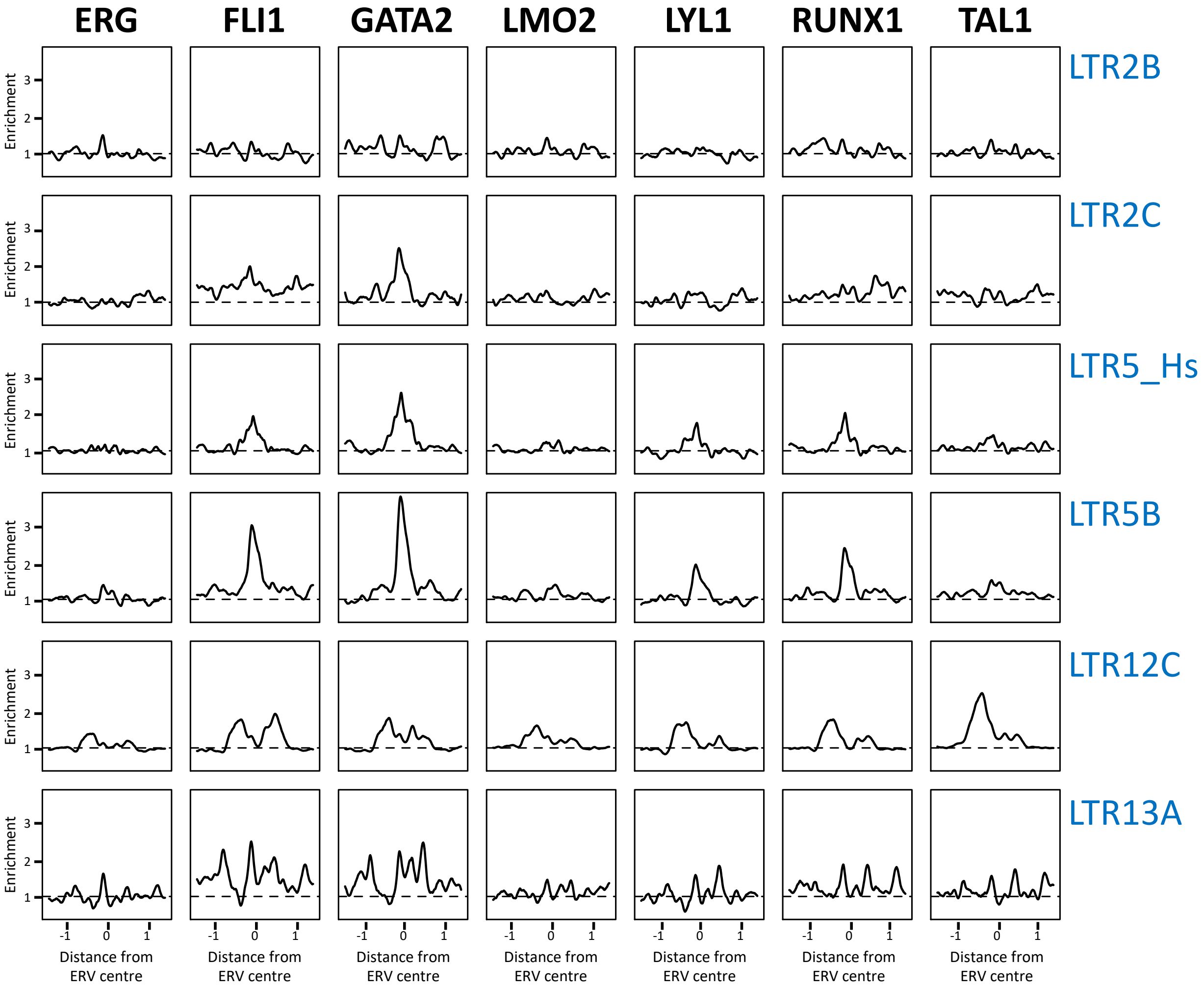


**Supplementary Figure 4.** Average ChIP-seq profiles of the given TFs for each A-DAR family in CD34+ hematopoietic progenitors. Data from the BloodChIP database.

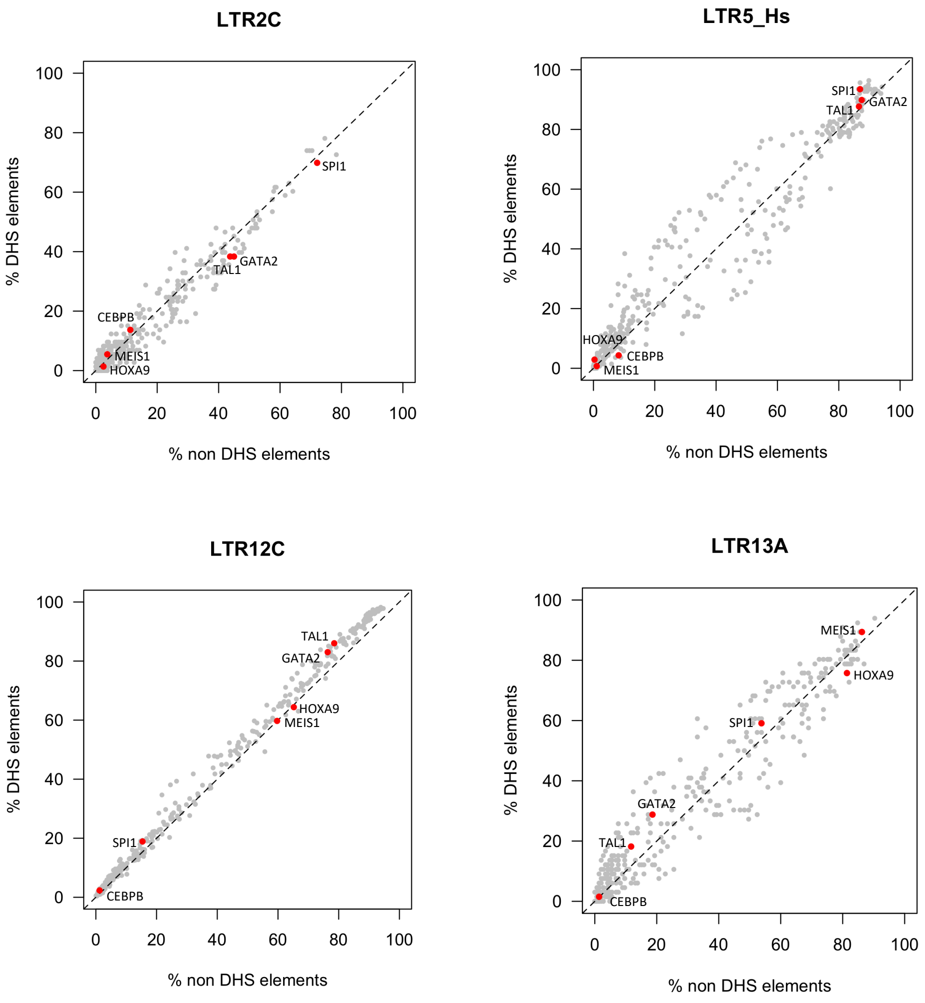


**Supplementary Figure 5.** TF motif frequency at LTR2C, LTR5_Hs, LTR12C and LTR13A elements, comparing those that overlap DHSs with those that do not.

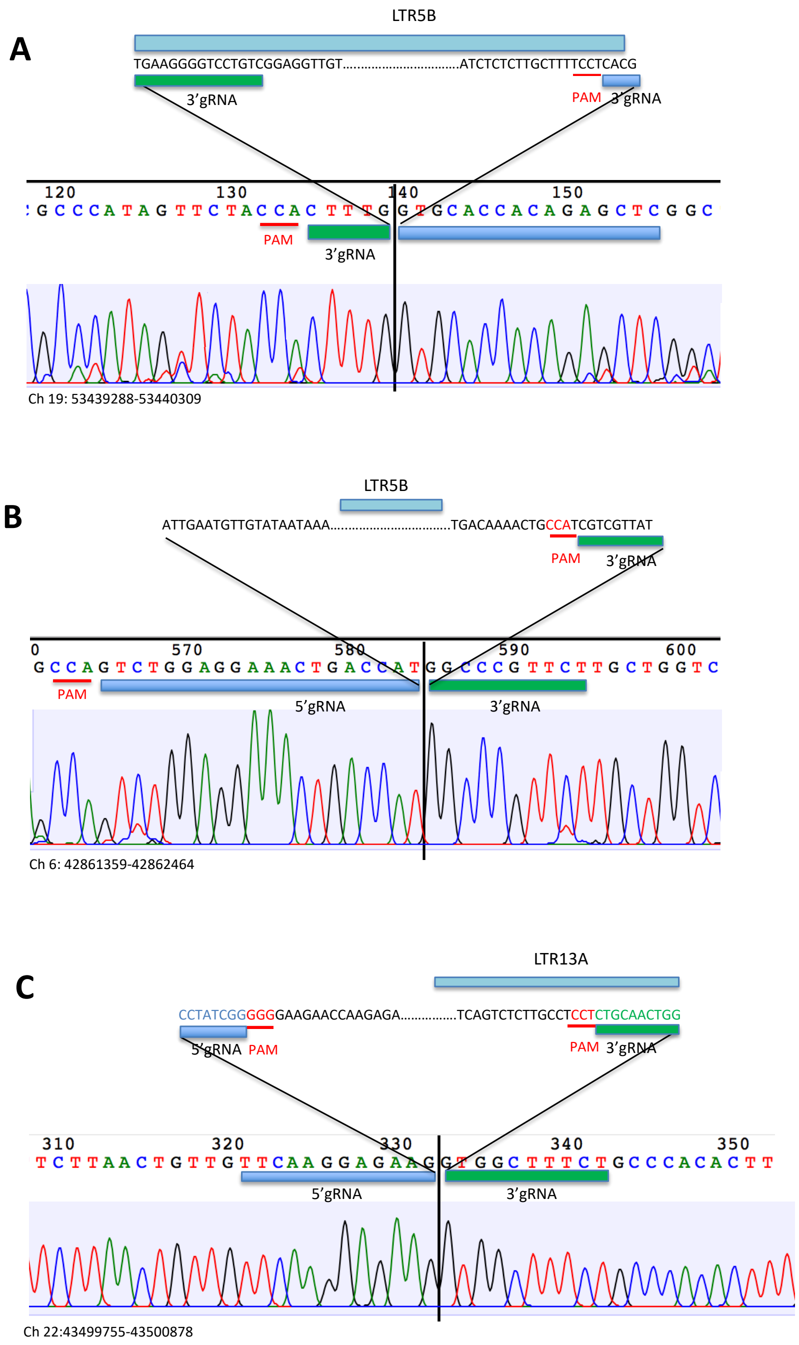


**Supplementary Figure 6.** Chromatography of the representative excision clones of three candidate regulatory ERVs. Sanger sequencing results are obtained from the alleles, where the indicated ERV is deleted.

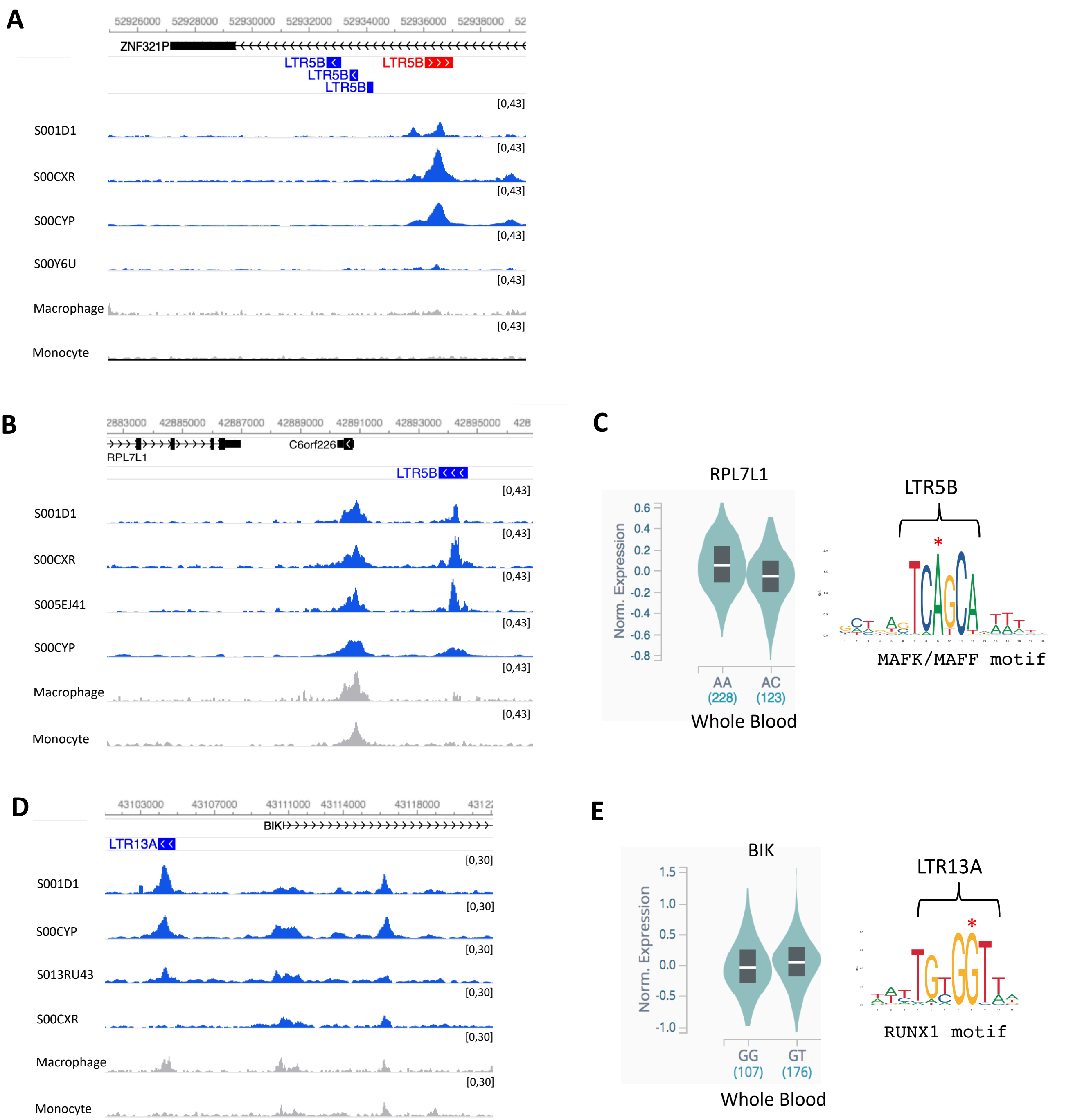


**Supplementary Figure 7. A.,B.,D.** Genome browser view of three candidate regulatory ERVs, showing DNase-seq tracks in AML, macrophage and monocyte samples. **C,E.** Location of SNPs within LTR5B (C) and LTR13A (D) (indicated by red asterix) are within a consensus MAFK/MAFF and RUNX1 binding motifs, respectively. Blood expression levels of RPL7L1 (C) and BIK (E) in heterozygous genotypes of the indicated SNPs.

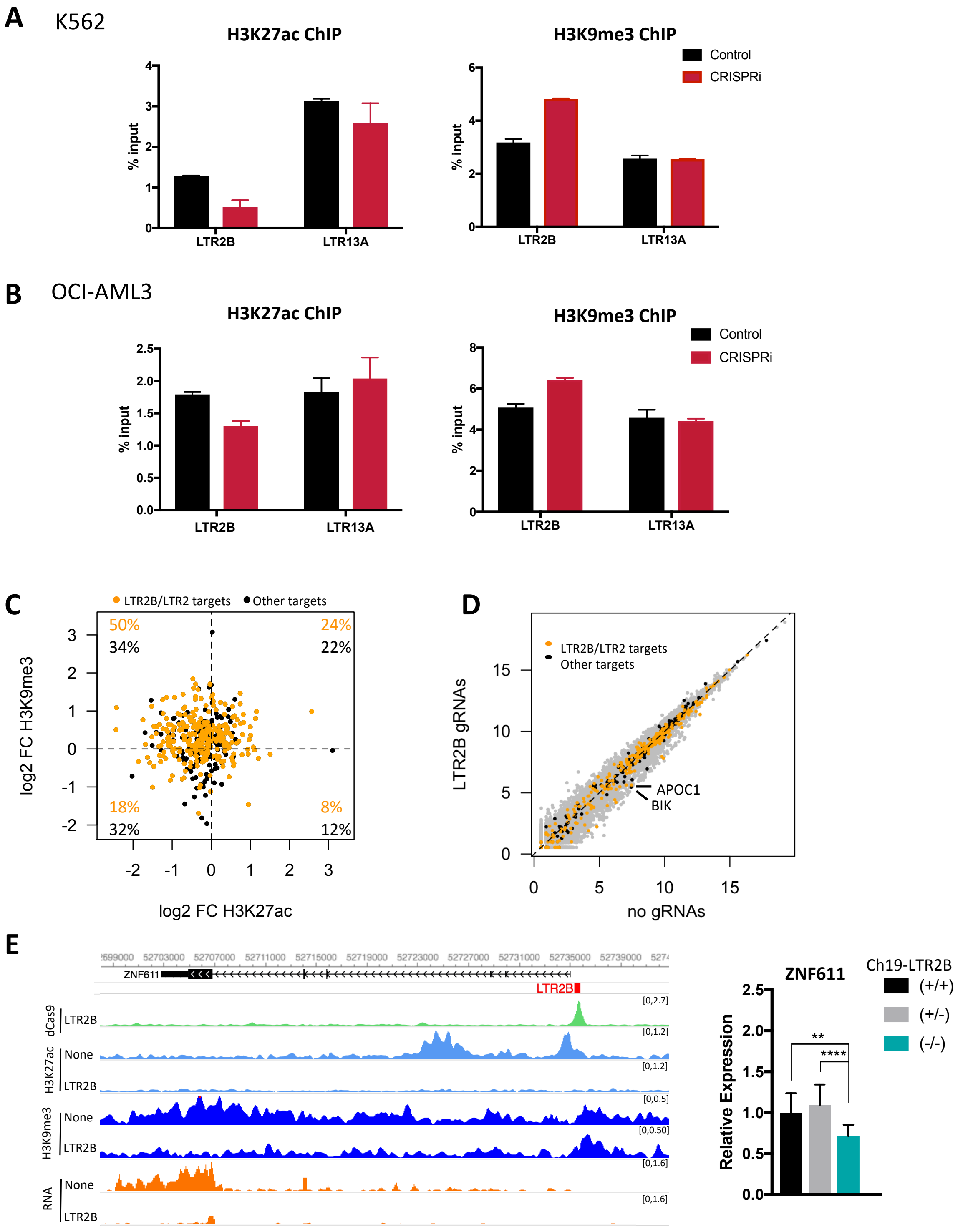


**Supplementary Figure 8. A.,B.** H3K27ac (left) and H3K9me (right) ChIP-qPCR at LTR2B elements upon CRISPRi in K562 (A) and in OCI-AML3 (B). **C.** Log2 ratio of the ChIP-seq signal at dCas9 peaks (1kb regions from the centre of each peak) between OCI-AML3 cells expressing LTR2B sgRNAs or empty vector. Orange points highlight dCas9 peaks overlapping LTR2B or LTR2 elements. **D.** Gene expression levels in OCI-AML3 cells expressing LTR2B sgRNAs or empty vector. Orange points highlight genes within 50kb of a dCas9 peak (in K562 cells) targeting LTR2B/LTR2 elements; black points refer to genes within 50kb of other dCas9 peaks. **E.** Genome browser snapshot for *ZNF611*-LTR2B element showing H3K27ac, H3K9me3 ChIP-seq and RNA-seq tracks in K562 cells expressing LTR2B sgRNAs or empty vector (left), expression of ZNF611 in the excision clones of *ZNF611*-LTR2B element. The error bars show standard deviation (n=8 technical replicates for each independent clone: 2 +/+, 4+/-, 2-/-, ANOVA with Tukey's multiple comparison test, $**p < 0.01$, $****p < 0.0001$).

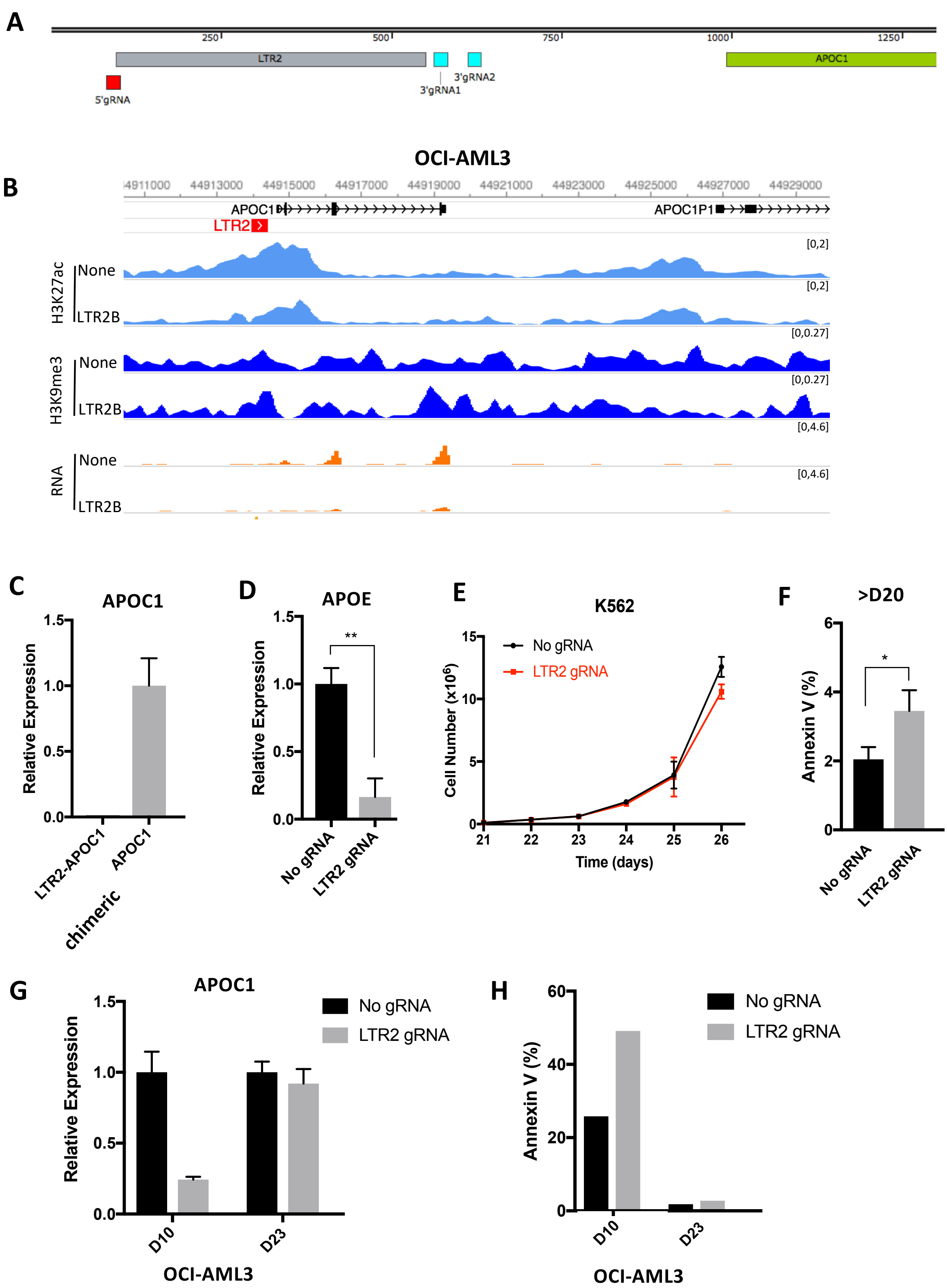


**Supplementary Figure 9**. **A.** The location of the designed gRNAs targeting LTR2-*APOC1.* **B.** Genome browser snapshot for *APOC1*-LTR2, showing H3K27ac, H3K9me3 ChIP-seq and RNA-seq tracks for control or CRISPRi OCI-AML3 cells. **C.** Expression of chimeric transcript (LTR2-*APOC1*) compared to *APOC1* transcript in K562 WT cells . The error bars show standard deviation (n=3 technical replicates). **D.** Expression of *APOE* at D6 in control and edited K562 cells. The error bars show standard deviation (n=3 biological replicates, n≥3 biological replicates, ANOVA with Tukey's multiple comparison test, **$p < 0.01$). **E.** Cell proliferation assay at day >20 in K562 cells expressing *APOC1*-LTR2 sgRNAs or empty vector (n=2 biological replicates). **F.** % of Annexin V stained cells at day >20. The error bars show standard deviation (n=3 biological replicates, t-test, *$p < 0.05$). **G.** Expression of *APOC1* at D10 and D23 in control and edited OCI-AML3 cells. The error bars show standard deviation (n=3 technical replicates). **H.** % of cells stained by Annexin V at D10 and D23 in OCI-AML3.

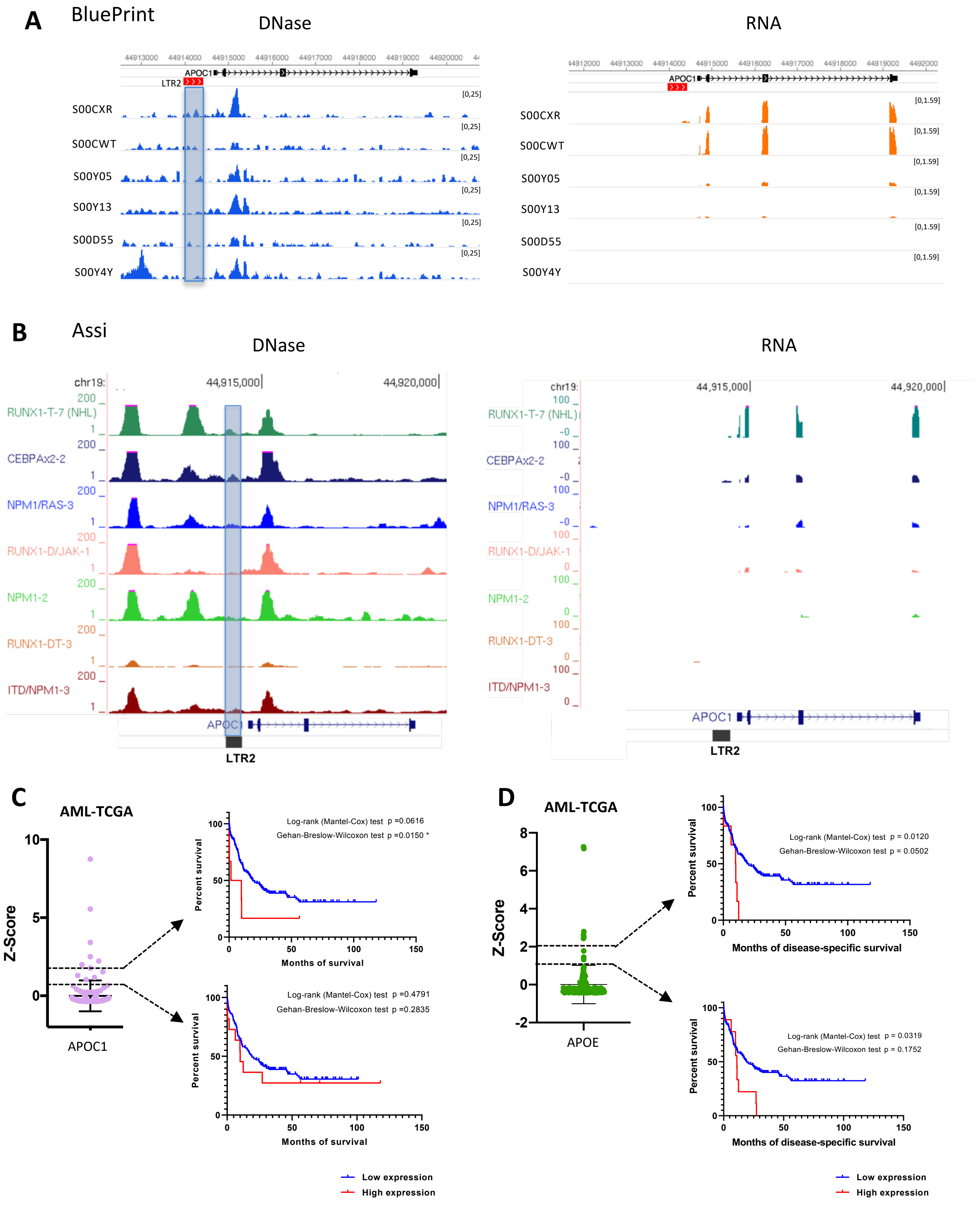


**Supplementary Figure 10. A,B.** Genome browser view of *APOC1*-LTR2, showing DNase-seq and RNA-seq tracks in AML samples from BluePrint (A) and Assi et al (B) datasets. **C,D.** Expression of *APOC1* (C) and *APOE* (D) based on The Cancer Genome Atlas (TCGA) (left) and Kaplan-Meier survival analysis according to *APOC1* (C) and *APOE* (D) expression (right) in AML. Results using two different expression cut-offs (indicated by the dashed lines) are presented to highlight that significant differences in survival are only observed when using a stringent threshold.
